# NMDA receptors require multiple pre-opening gating steps for efficient synaptic activity

**DOI:** 10.1101/2020.06.09.142687

**Authors:** Johansen Amin, Aaron Gochman, Miaomiao He, Noele Certain, Lonnie P. Wollmuth

## Abstract

NMDA receptors (NMDAR) are glutamate-gated ion channels that mediate the majority of fast excitatory synaptic transmission in the nervous system. A central feature of NMDAR physiology is the opening of the ion channel driven by presynaptically-released glutamate. Using glutamate applications to outside-out patches containing a single NMDAR in the continuous presence of the co-agonist glycine, we find that agonist-bound receptors transition to the open state via two conformations, an ‘unconstrained pre-active’ state that can rapidly transition to the open state and contributes to synaptic events, and a ‘constrained pre-active’ state that requires more energy and hence time to open and does not contribute to fast signaling. To define how agonist binding might drive these conformations, we decoupled the ligand-binding domains from specific transmembrane segments for the GluN1 and GluN2A subunits. Displacements of the central pore-forming M3 segments define the energy of fast channel opening. However, to enter the unconstrained conformation and contribute to fast signaling, a peripheral helix, the GluN2 pre-M1, must be displaced before the M3 segments move. This pre-M1 displacement is facilitated by the flexibility of another nearby peripheral element, the GluN1 and GluN2A S2-M4. We conclude that peripheral structural elements – pre-M1 and S2-M4 – work in concert to remove constraints and prime the channel for rapid opening, thus facilitating fast synaptic transmission.

## INTRODUCTION

The transition from the closed, non-conducting conformation to the open, conducting conformation is the central physiological function of ion channels. The preferred low-energy conformation is generally the closed state, and channels require the input of energy to transition to the open state (Hille, 2001). For voltage-gated ion channels, displacements of the voltage-sensing domains generate this energy (Bezanilla, 2008; Toombes & Swartz, 2014; Ahern *et al.*, 2016), whereas for ligand-gated ion channels, it arises from agonist-induced conformational changes (Auerbach, 2013; daCosta & Baenziger, 2013; Plested, 2016). High-resolution structures of the closed and open conformations provide the endpoints for ion channel opening, but do not reveal how energy is communicated between structural elements nor potential structural intermediates, which are typically extremely transient and energetically unstable. Indeed, almost all channels transition from the closed to the open state via multiple structural intermediates (Aldrich *et al.*, 1983; Howe *et al.*, 1991; Colquhoun & Lape, 2012; Goldschen-Ohm *et al.*, 2013). These structural or conformational intermediates are physiologically critical since they dictate the kinetics and probability of pore opening, and hence how an ion channel contributes to membrane physiology.

Ionotropic glutamate receptors (iGluRs) are ligand-gated ion channels that mediate the majority of fast synaptic transmission in the central nervous system. Glutamate can activate three types of iGluRs: AMPA (AMPAR), kainate, and NMDA (NMDAR) receptors. iGluRs are highly modular, layered proteins (Figure 1A). To open the ion channel pore, the ligand-binding domain (LBD) must energetically communicate with the pore-forming transmembrane-domain (TMD). While this might arise from direct physical interactions between domains, the most likely pathway for energetic coupling is through a set of short polypeptide linkers, the LBD-TMD linkers, which connect the LBD to specific transmembrane segments (Figures 1B & 1C) (Schmid *et al.*, 2007; Talukder *et al.*, 2010; Yelshanskaya *et al.*, 2017).

**Figure 1.**
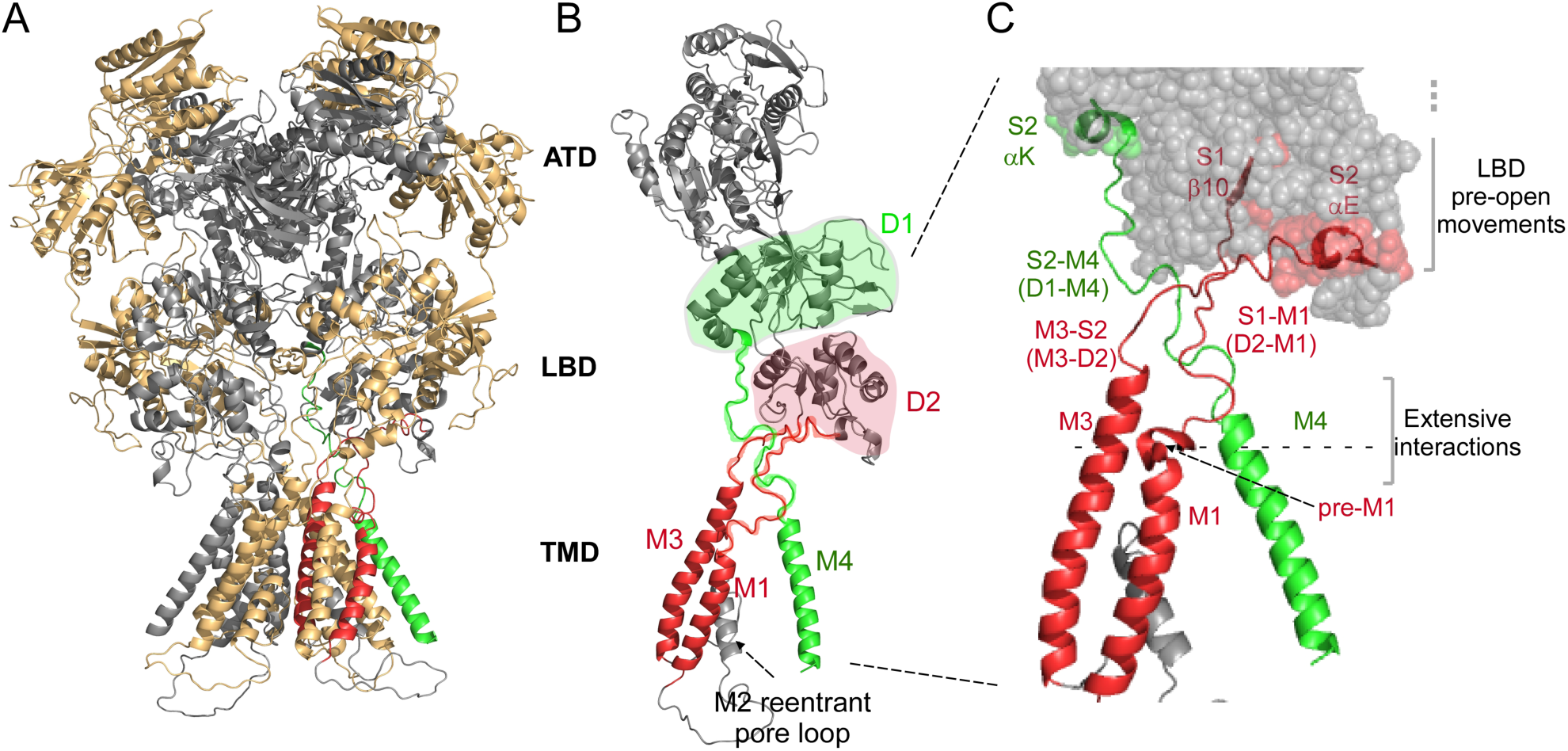
Structure and topology of NMDARs. **(A)** Ionotropic glutamate receptors (iGluRs) including NMDARs consist of four modular domains: the extracellular amino-terminal (ATD) and ligand-binding (LBD) domains; the transmembrane domain (TMD) forming the ion channel; and the intracellular C-terminal domain (CTD), which is absent in available high-resolution structures. NMDAR model structure based on 4TLM (Lee *et al.*, 2014; Amin *et al.*, 2017). Subunits are colored light orange (GluN1) and gray (GluN2B). **(B)** Topology of an individual subunit. LBD lobes are colored green (D1) or red (D2). The ion channel is formed by 3 transmembrane segments, M1, M3, and M4 and an intracellular M2 pore loop. Transmembrane segments are colored according to the LBD lobe to which they are connected. **(C)** Bridging the LBD and TMD in each subunit are three LBD-TMD linkers: S1-M1 (D2-M1), M3-S2 (D2-M3), and S2-M4 (D1-M4). The most membrane proximal secondary structures in the LBD are: β10 adjacent to S1-M1; αE adjacent to M3-S2; and αK adjacent to S2-M4. With agonist binding these structures undergo movements that precede ion channel opening (Tajima *et al.*, 2016; Zhu *et al.*, 2016; Chen *et al.*, 2017; Twomey *et al.*, 2017). At the top of the transmembrane segments, there are extensive interactions between the S1-M1/pre-M1/M1 and M3-S2/M3 of the same subunit and the S2-M4/M4 of an adjacent subunit. Pre-M1 is a short helix in S1-M1 that surrounds the corresponding M3 segment (Sobolevsky *et al.*, 2009).

In iGluRs, the LBD is comprised of two separate polypeptides, S1 and S2, which form a clamshell-like configuration comprised of two lobes, an upper D1 (mainly composed of S1) and a membrane-proximal D2 (mainly composed of S2) (Figure 1B)(Armstrong *et al.*, 1998; Furukawa *et al.*, 2005; Mayer, 2005). The core of the ion channel is the pore domain, consisting of transmembrane segments M1 and M3 and a cytoplasmic re-entrant M2 pore loop (Figure 1B). This pore domain is evolutionarily related to an inverted K^+^ channel (Wo & Oswald, 1995; Wood *et al.*, 1995). Notably, the linkers coupling M1 (S1-M1) and M3 (M3-S2) to the LBD are D2-linked (Figure 1C). In contrast, the eukaryotic-specific M4 transmembrane segments, which surround the pore domain, are D1-linked (S2-M4).

In iGluRs, agonist binding induces “clamshell closure”, in which the D2 lobe pulls away from the lipid bilayer (Sun *et al.*, 2002; Twomey *et al.*, 2017; Twomey & Sobolevsky, 2018). Analogous to K^+^ channels, the M3 transmembrane segments are the major pore-lining element, with their bundle helical crossing forming a gate (Chang & Kuo, 2008; Sobolevsky *et al.*, 2009). The basic mechanism opening this M3 gate presumably entails mechanical pulling (Sun *et al.*, 2002), with the agonist-induced displacements of the D2 lobe away from the membrane generating tension in the M3-S2 linker, providing the energy necessary for pore opening (Kazi *et al.*, 2014; Ladislav *et al.*, 2018).

The non-pore forming transmembrane segments, M1 and M4, are also displaced during gating and are critical for efficient pore opening (Kazi *et al.*, 2013; Chen *et al.*, 2017; Dolino *et al.*, 2017; Twomey *et al.*, 2017). The S1-M1 linker includes a pre-M1 helix, which surrounds the inner M3 segments and regulates pore opening (Ogden *et al.*, 2017; Gibb *et al.*, 2018; Amin *et al.*, 2020; McDaniel *et al.*, 2020). The S2-M4 linker/M4 segment also regulates gating (Amin *et al.*, 2017; Yelshanskaya *et al.*, 2017; Shi *et al.*, 2019). Still, the energetics and timing of these non-pore forming elements to receptor gating remains incomplete.

Here, we address the gating mechanism in NMDARs. At most synapses, NMDARs are obligate hetero-tetramers composed of two GluN1 and two GluN2 (A-D) subunits. To define conformational changes between agonist binding and pore opening, we rapidly applied glutamate to outside-out patches containing single NMDARs and assessed the time between when glutamate was first applied to when the channel first opened (‘latency to 1^st^ opening’, see Materials and Methods)(Aldrich *et al.*, 1983). With this approach, we identified that wild type GluN1/GluN2A NMDARs, in the transition from agonist-bound closed to the open state, transitions through one of two pre-active states: an ‘unconstrained’ state that can rapidly transition to the open state and contributes to fast synaptic signaling, and a ‘constrained’ state that slowly transitions to the open state, presumably due to higher energy requirements, and does not contribute to fast signaling. Further, by decoupling the LBD from the TMD for each specific LBD-TMD linker (S1-M1, M3-S2, and S2-M4, Figure 1B & 1C) (Niu *et al.*, 2004; Kazi *et al.*, 2014), we show that the non-pore forming linkers uniquely influence pore-opening by altering the stability of these pre-active conformations. Thus, our results highlight the temporal and coordinated movements from agonist binding to ion channel opening in NMDARs under non-equilibrium conditions, closely mimicking NMDAR activation at excitatory synapses.

## MATERIALS AND METHODS

### Mutagenesis and expression

Site-directed insertions were made in GluN1 (GluN1-a) (NCBI Protein database accession no. P35439) or GluN2A (Q00959) subunits with the QuikChange site-directed mutagenesis kit (Agilent) with XL1-Blue super-competent cells. All constructs were tested in rat orthologue. Residue numbering included the signal peptide (GluN1, 18 residues; GluN2A, 19 residues). In previous publications, we typically used numbering for the mature protein.

We transiently co-transfected cDNA constructs of GluN1 and GluN2A into mammalian human embryonic kidney 293 (HEK293) with a separate pEGFP-Cl vector (Clontech) at a ratio of 4.5:4.5:1 (GluN1/GluN2A/EGFP) using X-treme GENE HP (Roche) (for details on cell culture and transfection see Amin et al., 2017). Cells were grown at 37°C and 5% CO_2_ in Dulbecco’s modified Eagle’s medium (DMEM) containing 10% fetal bovine serum (FBS) for 24 h before transfection. To increase cell viability via limiting Ca^2+^ influx, the media bathing transfected cells also contained the NMDAR competitive antagonist APV (100 µM) and non-competitive antagonist Mg^2+^ (100 µM). Patch clamp recordings were performed 16-48 h post-transfection.

### Structural analysis

Analysis of available structures was done using PyMOL (http://pymol.org) for either NMDAR (PDBs 5fxh and 5fxg) (Tajima *et al.*, 2016) or AMPAR (5wek and 5weo) (Twomey *et al.*, 2017). To define a reference point for each set of structures, we identified the center of mass for the structure and would adjust slightly this reference point such that it was positioned near the center of the mouth for the ion channel pore, approximately just above the M3 helices bundle helical crossing. Using the measuring function in PyMOL, we measured from this reference point to the last αC in the designated secondary structure either in the closed (5wek) or open (5weo) state.

### Outside-out single channel recordings

Outside-out single channel recordings were collected at room temperature (21-24°C) using an EPC-10 patch clamp amplifier interfaced with PatchMaster (HEKA). We used thick-walled, borosilicate glass (Sutter Instrument) for patch pipettes, which were pulled and fire-polished yielding resistances between 5 and 20 MΩ when measured in the bath solution (with applied positive pressure of ∼150 mbar). Patch pipettes contained 140 mM KCl, 10 mM HEPES, and 1 mM BAPTA (pH 7.2, NaOH).

For all of the outside-out patch experiments, we used an external solution containing (in mM): 150 NaCl, 10 HEPES, 0.05 EDTA, pH 8.0 (NaOH). Although this external solution does not mimic what occurs at synapses, we used it since it removes the complications of proton (high pH) and divalent (Zn^2+^) inhibitory effects (Popescu & Auerbach, 2003) and the short- (Maki & Popescu, 2014) and long- (Legendre *et al.*, 1993) term effects of Ca^2+^ on NMDARs. We applied 1 s pulses of either 1 mM glutamate or 0.1 mM glycine through a piezo-driven double-barrel application pipette system (10-90% rise time of 400-600 µs). For both glutamate and glycine application experiments, the baseline barrel contained our standard Na-based external and either 0.1 mM glycine (for glutamate test applications) or 1 mM glutamate (for glycine test applications) while the test barrel contained the identical solution except for added 1 mM glutamate or 0.1 mM glycine, respectively. There were 4 s intervals in between pulses to allow recovery from desensitization (tau of recovery from desensitization for wild type GluN1/GluN2A is around 1 sec (Alsaloum *et al.*, 2016)). Currents were recorded at -70 mV.

For these experiments, we were mainly interested in the latency between when glutamate was first applied and the 1st opening of the ion channel (latency to 1^st^ opening). Given the high single channel current amplitudes of GluN1/GluN2A NMDARs which greatly facilitated detection of the 1^st^ opening, we did not rigorously control for noise. This allowed for a large number of patches/events to be recorded, greatly improving statistical analysis.

### On-cell single channel recordings

On-cell single channel recordings were collected at room temperature (21-24°C) on-cell single channel recordings of NMDARs at steady state, using an Axopatch 200B (Molecular Devices) integrating patch clamp amplifier. All on-cell patches were held at -100 mV. Currents were analog filtered at 10 kHz (lowpass Bessel filter) and digitized at 50 kHz (ITC-16 interfaced with PatchMaster, HEKA). The recordings were ∼4-60 min in duration to provide a substantial number of events for analysis. In general, we did not include on-cell patches in analysis unless it contained a minimum of 10,000 (typically >50,000) events, except for some of the GluN2A S1-M1 constructs that had extremely low open probability.

The bath (patch pipette) solution contained the same external solution as used for outside-out patches plus 1 mM glutamate and 0.1 mM glycine. Patch pipettes were the same as used in outside-out cell recordings and had resistances between 5 and 35 MΩ. Seal resistances were between 1 and 15 GΩ.

### Macroscopic or whole-cell currents

Whole-cell currents were recorded at room temperature (21-24°C) using an EPC-10 amplifier with Patchmaster software, digitized at 10 kHz and low-pass filtered at 2.9 kHz (−3 dB) using an 8 pole low pass Bessel filter (Yelshansky *et al.*, 2004). Patch microelectrodes were filled with an intracellular solution (in mM): 140 KCl, 10 HEPES, 1 BAPTA, pH 7.2 (KOH). Our standard extracellular solution consisted of (in mM): 140 NaCl, 1 CaCl_2_, 10 HEPES, pH 7.2 (NaOH). We recorded currents with or without 0.01 mM EDTA which was added to our standard extracellular solution to approximate conditions used in single channels. Pipettes had resistances of 2-6 MΩ when filled with the pipette solution and measured in the Na^+^ external solution. We did not use series resistance compensation, nor did we correct for junction potentials. Currents were measured within 15 minutes of going whole-cell.

### Analysis of single channel outside-out patches

Outside-out patches were exposed to super-saturating concentrations of glutamate (1 mM) and glycine (100 μM) (Traynelis *et al.*, 2010) to ensure receptor saturation and that the agonist binding steps occurred as rapidly as possible. At synapses, presynaptic-released glutamate appears rapidly, but is also present only transiently for ∼1 to 2 ms (Clements *et al.*, 1992). For our experiments, we applied glutamate for 1 second, which allowed us to readily determine whether patches contained just a single receptor, facilitating obtaining a high number of events. Given the high open probability and relatively long open time of most of the constructs tested, we are quite confident that patches contained a single receptor; many patches were rejected because of 2 or more channels. In contrast, several constructs, most notably glycine insertions in GluN2A S1-M1 and M3-S2, had low open probability and brief open times, making it less certain that patches contained a single channel.

To analyze outside-out patches, we exported data from PatchMaster to Igor, where they were analyzed using in-house developed programs. Applications displaying significant amounts of noise were removed (∼5–10% of applications). Nevertheless, noise was not rigorously controlled for since it was extremely straight-forward to detect the large NMDAR current amplitudes.

### Latency to 1^st^ opening

Durations from the start of an application to the first open event (latency or latency to 1^st^ opening) were reported. For each individual record, we estimated single channel amplitudes and used a threshold crossing to assist in identifying the 1^st^ opening. Each individual application was subsequently visually inspected to verify amplitude and latency time.

Latency to 1^st^ opening times were pooled and imported into ChannelLab (Synaptosoft). Latencies were binned at ∼ 60 μs intervals and histograms displaying number of events as a function of latency to 1^st^ opening were generated. The cumulative histogram was fitted by exponential components, until the log likelihood score could not be further improved.

### Failure to open

In some instances, agonist application showed no discernible NMDAR-mediated currents either during or after agonist removal (2 seconds total). These instances are referred to as failures. In the text, successes = 1 – failure rate.

### Analysis of single channel on-cell recordings

Analysis of single-channel records was comparable to Talukder and Wollmuth, 2011 (Talukder & Wollmuth, 2011). Briefly, PatchMaster data were exported to QuB. Each recording was visually inspected in its entirety for multiple simultaneous openings, signal-to-noise fluctuations, high-frequency artifacts, and baseline drifts. Further processing was done to eliminate occasional obvious brief noise spikes (e.g., high amplitude, stereotypical exponential-like decay, etc) by matching them to the level of adjacent events using the ‘erase’ function in QuB. Long periods of high noise were deleted with the remaining flanking segments separated as discontinuous segments. Baseline drift was corrected by resetting baseline to zero current levels. Processed data were idealized using the segmental k-means (SKM) algorithm in QuB with a dead time of 40 μs (Qin, 2004). Single-channel amplitude (pA), equilibrium open probability (Eq. *P*_O_), mean closed time (MCT), mean open time (MOT), and number of events were extracted. Eq. *P*_O_ is the fractional occupancy of the open states in the entire single-channel recording, including long lived closed states.

### Identification of single channel patches

For wild type GluN1/GluN2A and S2-M4 insertion constructs, the high *Po* allowed us to distinguish single-channels quite easily, as recordings with multiple channels had apparent multiple conductance states (Colquhoun & Hawkes, 1990). For low Eq. *Po* S1-M1 and M3-S2 glycine insertion constructs, we ensured that recordings were of long duration (∼10,000-150,000 events) to maximize the likelihood of observing multiple conductance states. Nevertheless, certain constructs showed an extremely low Eq. *Po* (<0.07). Although we did not detect multiple conductance states, we cannot rule out that these patches contained more than one channel.

### Analysis of macroscopic currents

To determine the extent and rate of desensitization in the whole-cell mode, we applied glutamate at -70 mV for 2.5 secs. Percent desensitization (% des) was calculated from the ratio of peak (I_peak_) and steady-state (I_ss_) current amplitudes: %des = 100 x (1 – I_ss_/I_peak_). Time constants of desensitization (weighted τ) were determined by fitting the decaying phase of currents to a double exponential function. In some instances when current amplitudes were small, we averaged 3-8 records.

To determine the rates of activation and deactivation in the whole-cell mode, we applied glutamate for 2 ms at -70 mV. For activation times, we used the 10-90% rise time. Deactivation times (weighted τs) were derived by fitting the decay phase of currents with a double-exponential function.

### Summed currents for outside-out patches

Where possible, currents were summed for outside-out patches. These summed currents were analyzed for peak and steady state current (to derive %des), rate of desensitization (weighted τ_des_), activation rate (10-90% rise time), and deactivation rate (fitted with double exponential function to derive weighted τ_deact_). In some instances, we were not able to analyze a particular feature of summed currents either because of too few events and/or too strong of desensitization.

### Thermodynamics analysis

We quantified the Gibbs free energy for the transition rate to activation:

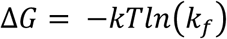

where k_f_ (= 1/τ) is the forward rate constant for activation, either for the fast or slow pathway. tau is derived from the fitted exponential.

For our experiments, we identified two kinetic components of the latency to 1^st^ opening. We generally focused quantitatively on the energetics and properties of the fast component. The specific energetics of the slow component would depend on the duration of the glutamate application (subsets of failures might be included with longer glutamate applications, which would alter the overall slow component distribution).

### Statistics

Data analysis was performed using IgorPro, QuB, Excel, and ChannelLab. Results from QuB or Igor analysis for each recording were organized in a Microsoft Excel sheet. All average values are presented as mean ± S.E.M. For statistical analysis, we used Excel or MiniTab 18. In instances where we were only interested in whether outcomes were statistically different from those of wild type, we used an unpaired two-tailed Student’s t-test to test for significant differences. Statistical significance was typically set at *p* < 0.05. In instances where we were interested in how constructs varied from each other, we used an ANOVA and followed with Tukey’s test (*p* < 0.05).

We did not run a statistical test to determine sample size *a priori*. Sample sizes resemble those from previous publications.

## RESULTS

To address the coupling between agonist-induced displacements of the ligand-binding domain (LBD) and ion channel opening, we recorded from outside-out patches containing a single NMDAR and rapidly applied glutamate (Figure 2A). As presumably occurs at synapses, channels were continuously bathed in glycine. We focused on the time interval between when glutamate was initially applied to the first identifiable opening of the ion channel. We refer to this period as the ‘latency to 1^st^ opening’ (Aldrich *et al.*, 1983). This latency has physiological significance since it reflects how efficiently a glutamate signal is translated into ion channel opening. In addition, latency to 1^st^ opening is the most direct assay of the coupling between LBD movements and those of the transmembrane segments leading to pore opening since it requires no assumptions about idealization and simultaneously captures non-equilibrium conditions for a single channel.

**Figure 2.**
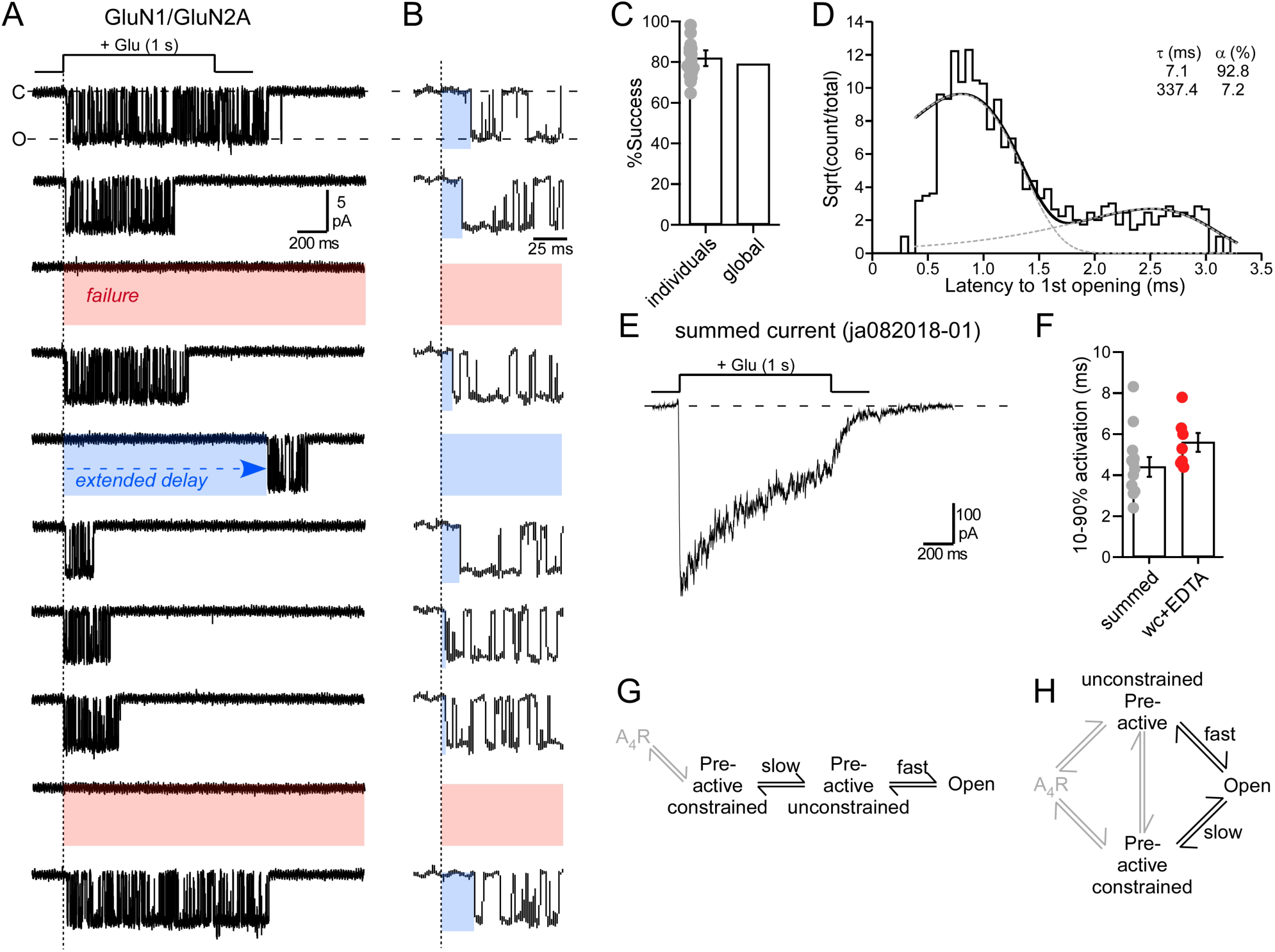
Agonist application to single channel outside-out patches containing wild type GluN1/GluN2A. (**A & B**) Representative current traces from an outside-out single channel patch (10 consecutive traces from same recording) of wild-type GluN1/GluN2A. Traces in (**B**) are the same as those in (**A**) but on an expanded time scale. The patch was exposed to pulses (1 s) of glutamate (1 mM) in the continuous presence of glycine (0.1 mM). Current responses reflected either successes (channel opening) or failures (red highlight), with successes showing either a brief or an extended delay (trace 5) to first opening (blue highlight). Currents were sampled at 50 kHz (displayed at ∼ 1 kHz). Holding potential, -70 mV. **(C)** Success rate (1 – failure rate) for the average of all patches (18 total patches) or the global success rate (1279 total trials with 1011 successes). In this and all subsequent figures, dots represent individual data points. **(D)** Dwell time histogram of the latency to 1^st^ opening, the time between the start of agonist application and the 1^st^ ion channel opening. The histogram was best fit by 2 exponentials (dashed lines). **(E)** Example summed current. This patch had 140 (123 successes) applications. **(F)** Bar graph (mean ± SEM) showing activation rates (10-90% rise time) for summed currents (13 patches) from outside-out patches or from current recorded in the whole-cell (wc) mode (Supplemental Table 1). Values were not significantly different (*p = 0.069, t-test*). (**G & H**) Potential pathways for the transition of receptor to the open state. A_4_R is the receptor fully bound with agonist, which we assume occurs rapidly after the start of the glutamate application because we used a supersaturating concentration.

### Latency to 1^st^ openings in wild-type GluN1/GluN2A receptors

Wild-type GluN1/GluN2A channels typically activated shortly after glutamate exposure (Figure 2A, trace 1 & 2), but with variations (Figure 2B), including in some instances extended delays (trace 5). There were also instances of failures to open (3^rd^ & 9^th^ traces). For wild-type GluN1/GluN2A, we recorded from 18 different outside-out patches with the number of applications varying from 10 to up to 325 per patch. The latency to 1^st^ opening and success rate (=1 – failures) showed no time dependence (Supplemental Figure 1), indicating that they could be combined across the patches. We made a total of 1279 applications to the 18 patches of which 268 were failures, that is no single channel activation occurred over the tested time interval. The success rate of the 18 individual patches was 82.0 ± 2% (n = 18) (mean ± SEM, n = number single channel patches), whereas the global success rate was 79% (n = 1011, where n = number of successful events out of 1279 applications) (Figure 2C).

For successes, the mean latency to 1^st^ opening was 48.0 ± 7.9 ms (n = 18) (global mean latency, 47 ms, n = 1011). To characterize these latencies, we generated dwell time histograms (Figure 2D). The latency to 1^st^ opening dwell time histogram was best fit by 2 exponentials: a fast (τ = 7.1 ms, 92.8%) and a slow (τ = 337.4 ms, 7.2%) component. Thus, of the total successful events (1011), approximately 938 events corresponded to the fast component, while 73 events corresponded to the slow component. As a fraction of total trials (1279), 73.3% corresponded to the fast component, while 5.7% corresponded to the slow component (note that the remaining events were failures).

The fast component events would most likely contribute to synaptic events, whereas slow component events (and failures) would not. Hence, for wild type GluN1/GluN2A, synaptic efficiency – those fast component events out of the total number of trials – would be around 73% (see Discussion). Energetically, the activation energy through the fast component requires 1.96kT, whereas the slow component pathway requires 5.82kT (see Materials & Methods).

### Properties of summed currents

To further define the properties of the currents arising from these single channel patches, we summed these currents for each individual patch (e.g., Figure 2E) and characterized their properties (Supplemental Table 1). The activation rate of these summed currents (10-90% rise time, 4.4 ± 0.5 ms, n = 13) was comparable albeit somewhat faster than whole-cell recordings under comparable conditions (5.8 ± 0.5, n = 6) (Figures 2F; Supplemental Table 1), supporting the idea that the fast component could contribute to synaptic events.

One possibility to account for failures is that receptors are in a desensitized state when glutamate is applied, either because the receptors have not recovered from desensitization (see Materials & Methods) or because one of the LBDs is closed intrinsically (Yao *et al.*, 2013). Although we cannot rule out that desensitization contributes to failure rates, we do not believe it is a major factor since there is a delay between applications (see Materials & Methods) and there is no relationship (r^2^ = 0.09) between extent of desensitization and failure rates for individual patches (Supplemental Figure 2). Still, even if desensitization contributed to failure rates, it would not alter subsequent conclusions.

### Kinetic models of GluN1/GluN2A activation

Given the two kinetic components, we assume there are two agonist-bound closed states between glutamate binding and ion channel opening. We will refer to these states as ‘unconstrained pre-active’ associated with the fast component, and ‘constrained pre-active’ associated with the slow component (and also potentially failures)(see Materials & Methods). These two states could be arranged linearly (Figure 2G) or in parallel (Figure 2H). Subsequent experiments will favor the parallel pathway (Figure 2H) (see Discussion). In this context and in terms of the physiology of NMDARs, there are two critical considerations: (i) The availability of receptors in the ‘unconstrained pre-active’ state, that permits receptors to transition rapidly into an open conformation and contribute to synaptic signaling. We refer this feature as ‘efficiency’. (ii) The amount of energy required to open the channel in the ‘fast’ pathway. Hence, in the ionic conditions used in the present set of experiments, 73% of the receptor exist in the more efficient ‘unconstrained pre-active’ state and it takes 1.96kT to open the channel.

### D1- and D2-associated linkers show different displacements with clamshell closure

To begin to understand the structural basis for the coupling between the LBD and specific transmembrane elements, we quantified the displacement of the most membrane proximal secondary structure in the LBD (αE, β10, & αK)(Figure 1C) in the transition from the clam-shell open (ion channel closed) to the clam-shelled closed (ion channel open) conformations with αE coupled to M3 via M3-S2 (Figures 3A & 3B); β10 coupled to pre-M1/M1 via S1-M1 (Figure 3C & 3D), and αK coupled to M4 via S2-M4 (Figures 3E & 3F). For this analysis, we used the AMPAR closed (5wek) and open (5weo) state structures (Twomey *et al.*, 2017), because the end states are better defined and the relationship of the LBD-TMD linkers could be visualized. Although potentially comparable conformations exist for NMDARs (Tajima *et al.*, 2016; Zhu *et al.*, 2016), there is ambiguity as to the conformation of the presumed pre-active states.

**Figure 3.**
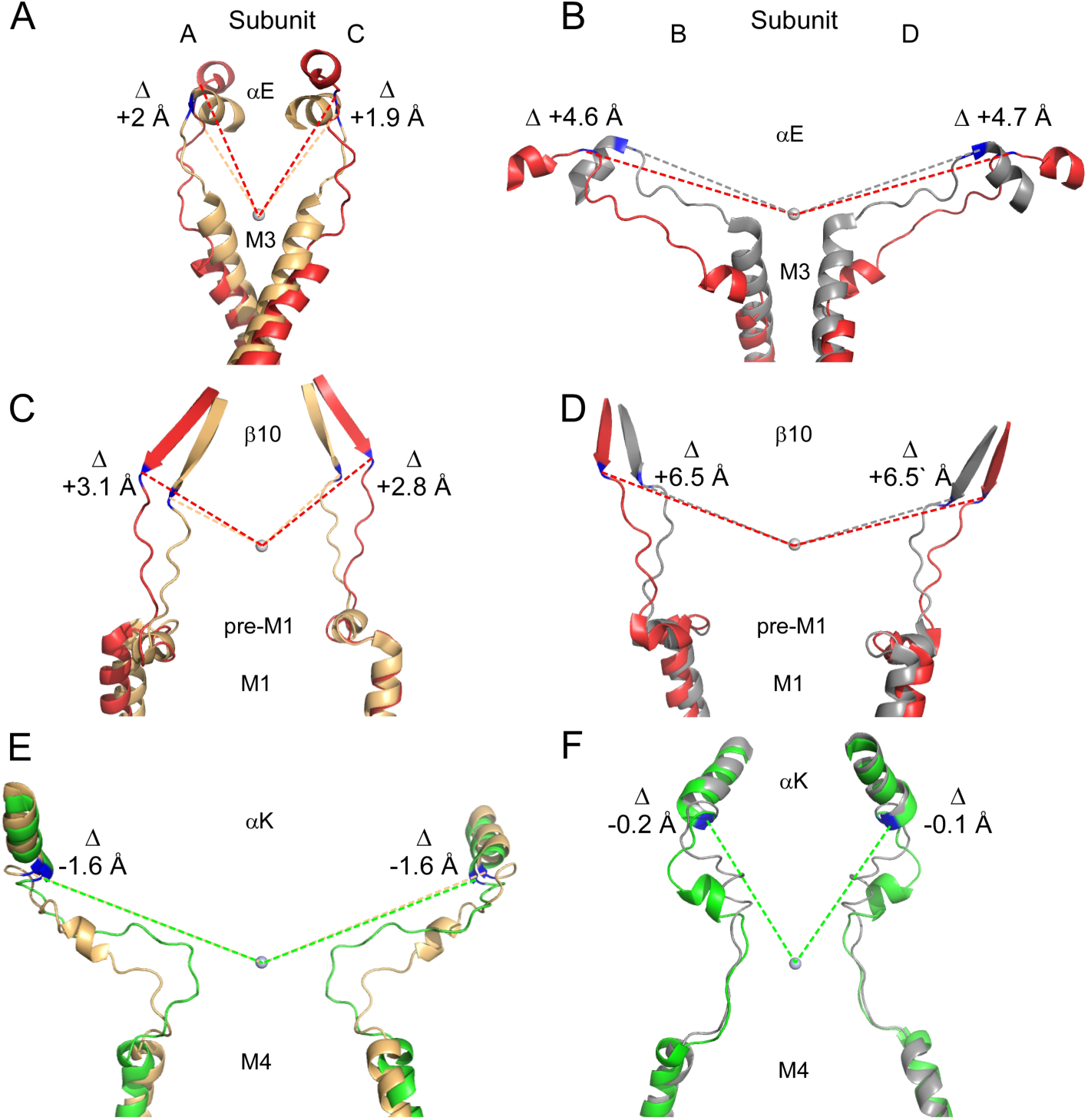
Displacement of membrane proximal secondary structures in the transition from clamshell open to clamshell closed. (**A-F**) The most membrane proximal structures to the LBD-TMD linkers either move away (D2-associated) or toward (D1-associated) the M3 gate. Overlaid AMPAR structures in the closed (glutamate unbound, 5wek) and open (glutamate bound, 5weo) conformations (Twomey *et al.*, 2017). Structures are in the A/C (∼GluN1) (**A, C, E)** or B/D (∼GluN2A) (**B, D, F**) conformations. In the A/C conformations, the closed state is light gold, whereas in B/D it is grey. The open state is red (D2-associated) or green (D1-associated). For quantification, we measured the distance from the reference point, the circle at the mouth of the pore (see Materials & Methods), to the last αC in the most membrane proximal secondary structure in the LBD: (**A & B**) αE coupled to M3 via M3-S2; (**C & D**) β10 coupled to pre-M1 via S1-M1; and (**E & F**) αK coupled to M4 via S2-M4. The numbers shown for each subunit are the difference between the open and closed state distance.

For this analysis, we distinguished the secondary structures into D2 (αE and β10), where the open state was highlighted in red (Figures 3A-3D), or D1 (αK), where the open state was highlighted in green (Figures 3E & 3F). Notably, although they take on distinct A/C (∼GluN1) and B/D (∼GluN2A) conformations (Salussolia *et al.*, 2011; Karakas & Furukawa, 2014; Lee *et al.*, 2014), all D2-associated secondary structures show a strong displacement away from the central axis of the pore with these displacements strongest in the B/D subunits. For αKs, the last αCs in A/C were displaced by +2 and +1.9 Å (Figure 3A), whereas for B/D they were +4.6 and +4.7 Å (Figure 3B), as would be expected for mechanical pulling. A similar general pattern is seen in β10, where the last αCs are displaced by +3.1 and +2.8 Å in A/C (Figure 3C) and by +6.5 and +6.5 Å in B/D (Figure 3D). In contrast, αKs which are part of D1 and linked to S2-M4 show a notable displacement towards the central axis of the pore: the last αCs are displaced by - 1.6 and -1.6 Å in A/C (Figure 3E), whereas by -0.2 and -0.1 Å in B/D (Figure 3A).

Hence, D2-associated structures (β10 & αE) show strong movements away from the M3 gate, consistent with some sort of mechanical pulling. On the other hand, D1-associated structures (αK) show a notable movement towards the membrane, which has unknown mechanistic significance.

### Decoupling the LBD from the TMD

Based on the structural analyses, we hypothesized that D1- and D2-associated linkers play different roles in NMDAR pore opening. To address the energetic coupling between agonist-induced displacements of the LBD and those of the transmembrane elements leading to pore opening, we inserted glycine residues at different points in the LBD-TMD linkers in the GluN1 and GluN2A subunits and subsequently quantified channel function (Kazi *et al.*, 2014). Our goal with these insertions was to increase the distance and/or flexibility between the LBD and transmembrane elements and hence decrease the efficiency of energy transfer from the LBD to the transmembrane elements. Our assumption is that such glycine insertions would mainly decouple the pre-open movements of a specific LBD element (the last secondary structure in the LBD) to its associated transmembrane element (Figure 1C). Insertions should have minimal effects on local interactions, which in themselves might change the energetics of channel opening. Glycine insertions, when tested, have no strong effect on the LBD dynamics (Kazi *et al.*, 2014).

### LBD coupling to the M3 segments determines the energy of the fast pore opening step

As an initial step to define energetic coupling, we looked at glycine insertions in the M3-S2 linkers (Figure 4; Supplemental Table 2)(Kazi *et al.*, 2014). The M3-S2 linkers couple αE in the D2 lobe of the LBD to the M3 segments (Figures 1C & 3). The M3 segments are the main pore lining segments and at their apex form an activation gate for ion channel opening (Chang & Kuo, 2008; Sobolevsky *et al.*, 2009). For these experiments, we used glycine insertions in either GluN1 (G648+1G) or GluN2A (G645+1G) M3-S2s that decouple the LBD from the M3 segment (Kazi *et al.*, 2014).

**Figure 4.**
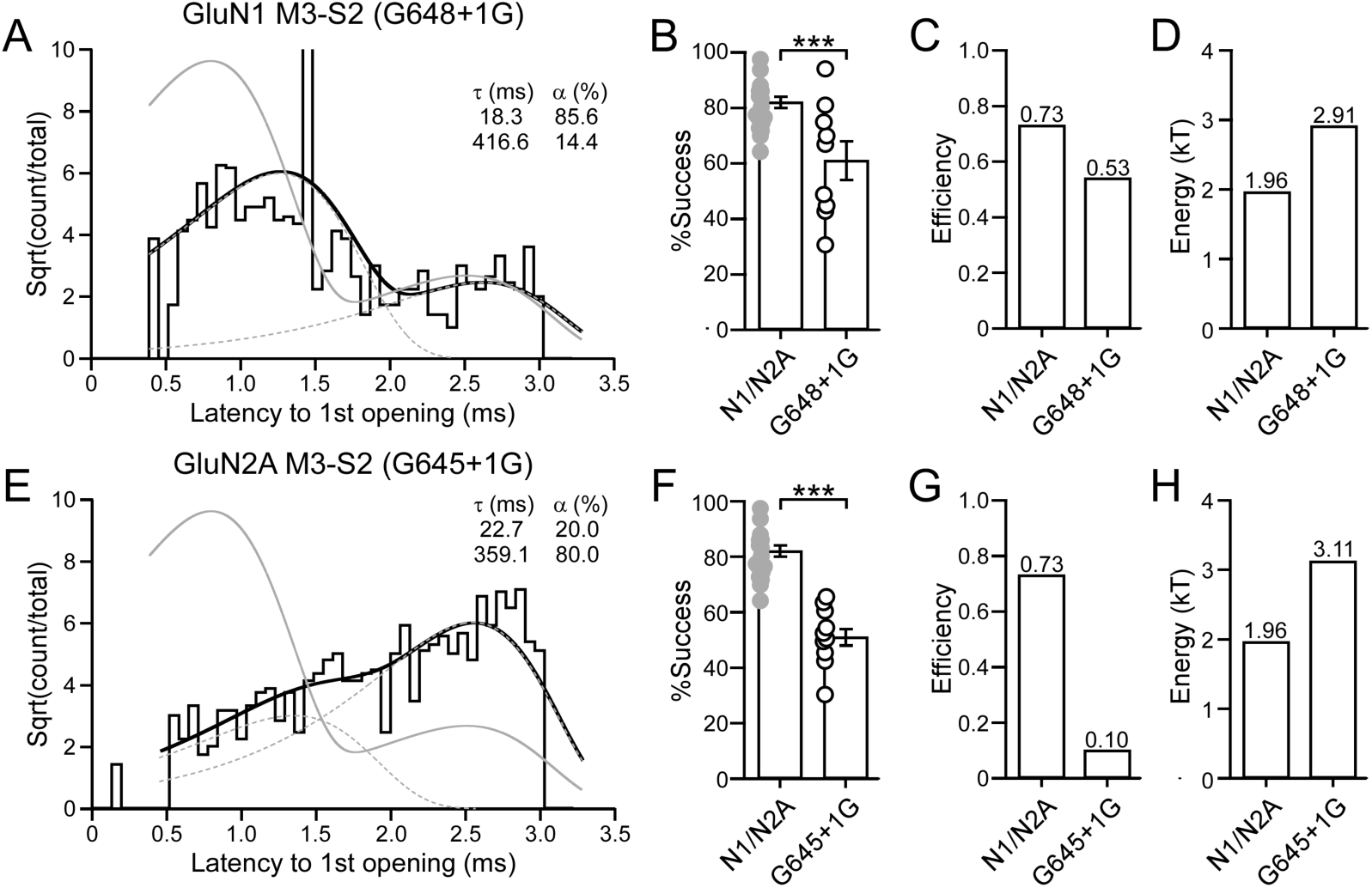
Glycine insertions in the GluN1 or GluN2A M3-S2 show two kinetic components. **(A)** Dwell time histogram of the latency to 1^st^ opening for a glycine insertion in GluN1 M3-S2 (G648+1G). The histogram was best fit by 2 exponentials (dashed lines). Grey line is the summed exponential fit for wild type. **(B)** Success rate (1 – failure rate) for the average of all patches for wild type or GluN1 M3-S2 +1G (9 total patches) (global success rate, 0.62; 315 out of 505 trials). *p* < *0.001, two-tailed Student’s t-test, unpaired*. **(C)** Efficiency of opening to the fast component: efficiency = (successes x fraction fast component)/total number of trials. **(D)** Energy requirement for fast component. **(E-H)** Same as A-D except for a glycine insertion in the GluN2A M3-S2 (G645+1G). (**F**) Success rate for GluN2A M3-S2 +1G (10 total patches) (global success rate, 0.51; 415 out of 808 trials). *p* < *0.001, two-tailed Student’s t-test, unpaired*. Note a subset of the data shown in Figure A & E were also shown in Kazi et al., 2014. However, we include here because of the distinct analysis.

Notably, fits to the dwell time histogram for the latency to 1^st^ opening for glycine insertions in either GluN1 (Figure 4A) or GluN2A (Figure 4E) M3-S2 required two kinetic components. Hence, transitions to a ‘fast’ and a ‘slow’ pathway still existed as in wild type NMDARs. Nevertheless, these components were shifted relative to wild type. In addition, glycine insertions in GluN1 (Figure 4B) and GluN2A (Figure 4F) M3-S2 showed significantly reduced successes.

For the GluN1 M3-S2+1G, the efficiency of the fast component for GluN1 was reduced from 0.73 to 0.53 (Figure 4C), reflecting both a reduction in the occupancy of the fast component as well as increased failures. Hence, about 53% of the time receptors existed in a conformation where they could transition ‘rapidly’ into an open state. On the other hand, 2.91kT (versus 1.96kT for wild type) of energy was needed to open the channel in the ‘fast’pathway (Figure 4D), presumably reflecting inefficient energy transfer from the LBD to M3. The situation was even more severe in the GluN2A subunit where the glycine insertion reduced efficiency to 0.10 (Figure 4G) and the energy required for the rapid transition was 3.11kT (Figure 4H).

In summary, the reduced entry into the fast component could arise in one of two ways: either the receptor in the absence of glutamate occupies the ‘unconstrained pre-active’ state only about 53% (GluN1) or 10% (GluN2A) of the time; alternatively, given the reduced energy transfer to the M3 segment, the energy generated in the M3 segments could be insufficient to overcome other structural constraints. Importantly, a greater amount of energy transfer is required to open the channel even when in the ‘fast’ pathway (Figures 4D & 4H). Hence, the mechanically pulling entailing the M3-S2 is directly involved in opening the ion channel, consistent with its central role in iGluR gating (Figures 3A & 3B)(Kazi *et al.*, 2014; Ladislav *et al.*, 2018).

### Glycine insertions in the GluN1 or GluN2A S1-M1 linkers attenuate receptor gating

In terms of the S1-M1, the most proximal secondary structure in the LBD is D2-associated β10 (Figures 3C & 3D, 5A & 5D) (Supplemental Figures 3 & 4). A notable feature of the S1-M1 linkers is the presence of pre-M1 helices that form a ring around the inner M3 segments (Sobolevsky *et al.*, 2009; Karakas & Furukawa, 2014; Lee *et al.*, 2014). These pre-M1 helices, especially those in the GluN2 subunit, play central roles in regulating the opening of the ion channel (Ogden *et al.*, 2017; McDaniel *et al.*, 2020).

We inserted glycines, individually, at largely equivalent points in the GluN1 (Figure 5A) and GluN2A (Figure 5D) S1-M1 linkers. The membrane proximal part of the S1-M1s, encompassing the pre-M1 helices (highlighted in grey in primary sequence), have many local interacting residues in the closed state (red arrows), whereas the remainder of the S1-M1s have fewer local interactions (blue arrows). Notably, there is a membrane-proximal proline (P) conserved across all mammalian iGluR subunits (Supplement Figures 3B & 4B) [GluN1(P557) & GluN2A(P552)] as well as an additional proline in GluN1 (P547) (Alsaloum *et al.*, 2016). Given the rigid secondary structure of proline, we inserted glycines on either side of these prolines. Numerous disease-associated missense mutations have been identified at the membrane proximal prolines (Ogden *et al.*, 2017; Soto *et al.*, 2019). To assay receptor function, we initially recorded the activity of single channel patches in the on-cell mode (See Materials & Methods).

**Figure 5.**
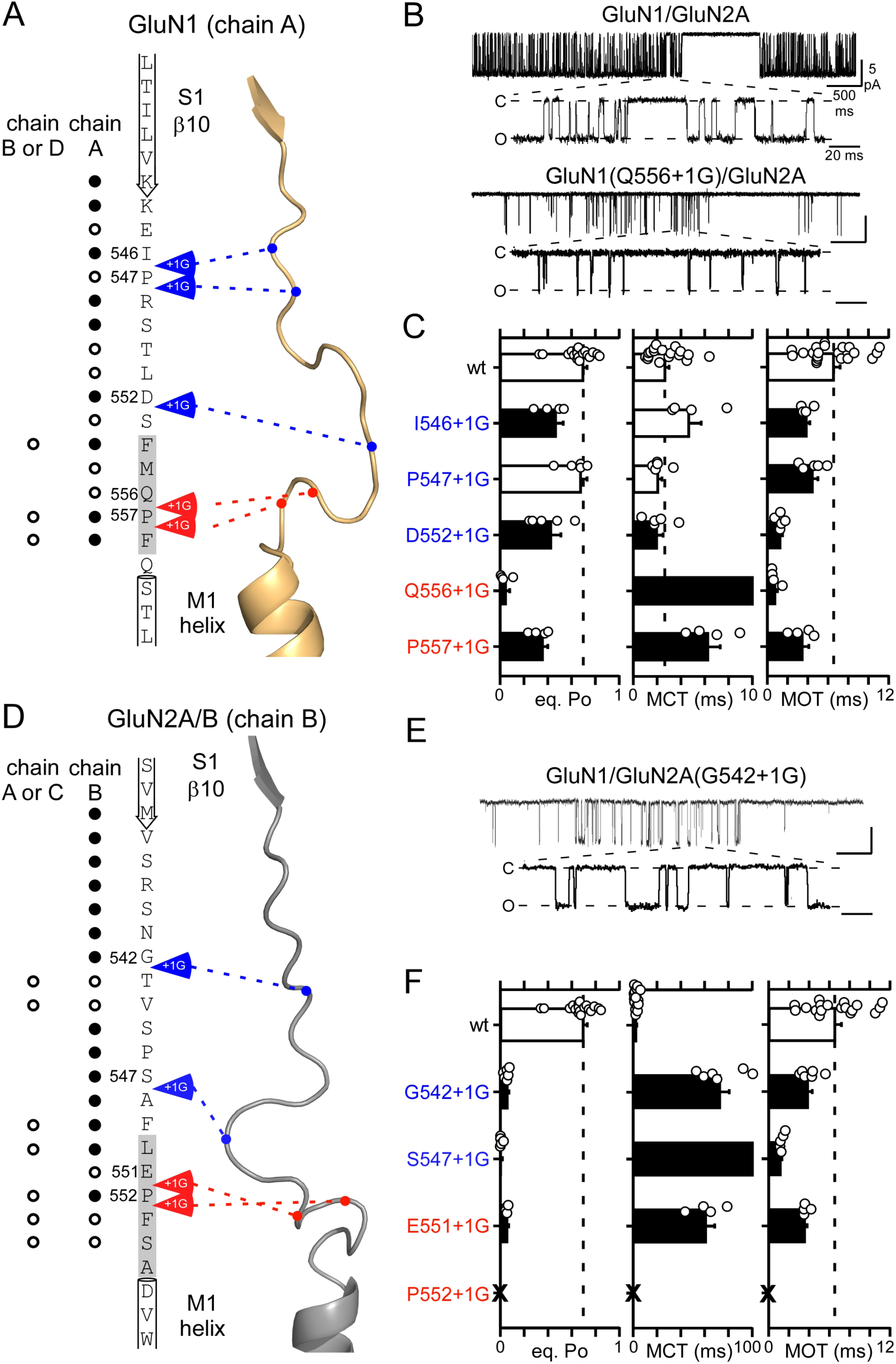
Glycine insertions in the GluN1 or GluN2A S1-M1 linkers reduce NMDAR gating. **(A & D)** Primary amino acid sequences and tertiary structures of the GluN1 (**A**) and GluN2B (**D**) S1-M1 linkers as well as the last structure in D2 (β10) and first three residues of the M1 segments. Grey shading in primary sequence highlight pre-M1 helices. Dots indicate the number of side chains in the closed state within 5 Å of that position [open circle, <5; solid circle, 5 or greater]. Locations of glycine insertions are indicated by +1G arrows along with amino acid numbers: Insertions not proximal to other structural elements are in blue, whereas those proximal are in red. Structural information is from 4TLM computational model (Supplemental Figures 3 & 4)(Amin *et al.*, 2017). **(B & E)** Representative traces from on-cell single-channel patches obtained at a -100 mV holding potential (downward inflections indicate inward currents). Experiments were conducted for wild-type GluN1/GluN2A and constructs containing single glycine insertions either in GluN1 (**B**) or GluN2A (**E**) S1-M1 linkers. In each panel, the upper trace captures five seconds of the recording, filtered at 1 kHz, while the lower trace in the expanded section is filtered at 4 kHz. Closed (C) and open (O) states are indicated in the higher resolution traces. Values for scale bars are shown in wild-type trace. **(C & F)** Single channel equilibrium open probabilities (*eq. P*_*o*_), mean closed times (MCT), and mean open times (MOT) (mean ± SEM) for wild-type and S1-M1 glycine insertion constructs either in GluN1 (**C**) or GluN2A (**F**). Dashed lines indicate mean wild-type values. Open circles are individual values. Solid bars indicate values significantly different from wild type (*p* < *0.05, two-tailed, unpaired Student’s t-test*). No single channel currents, indicated by ‘X’, could be detected for GluN2A(P552 +1G) as has been observed for substitutions at P552 (Ogden *et al.*, 2017). MCT values for Q556+1G and S547+1G are incompletely shown (see Supplemental Table 3).

Wild type NMDARs composed of GluN1/GluN2A and under the ionic conditions used in the present experiment (Figure 5B) show an equilibrium open probability (*eq. Po*) of around 0.70 (0.70 ± 0.03, n = 18)(mean ± SEM, n = number of independent on-cell patches) (Figure 5C)(Supplemental Table 3). With one exception, GluN1(P552+1G), all glycine insertions in the GluN1 (Figure 5C, *left panel*) and GluN2A (Figure 5F, *left panel*) S1-M1s significantly attenuated (black bars) single channel activity (Supplemental Table 3). These effects occurred whether the insertions were made membrane proximal or membrane distal.

There are four key observations from these glycine insertions in the S1-M1s. First, in general, the glycine insertions have a much stronger effect in the GluN2A S1-M1 than in the GluN1 S1-M1. Indeed, the *eq. Po* of the insertions in the GluN2A was considerably less than 0.1 whereas this occurred only for GluN1(Q556+1G) (Figures 5C & 5F, *left panel*; Supplemental Table 3). Second, the most consistent effect of the insertions independent of subunit was a significant reduction in mean open time (MOT) (Figures 5C & 5F, *right panel*). Third, while the insertions in GluN1 and GluN2A had about the same effect on reducing MOT, the stronger effect on eq. Po in the GluN2A subunits reflects that they had a much stronger effect on mean closed time (MCT) (Figures 5C & 5F, *middle panel*; note the changed scale in Figure 5F). Finally, the glycine insertions around the conserved membrane-proximal proline, as with disease-associated missense mutations at this proline (Ogden *et al.*, 2017), strongly inhibited receptor gating.

In summary, a presumed decoupling of the LBD (β10) from the pre-M1 helices strongly inhibits receptor gating. In addition, there is a stronger effect on GluN2A than on GluN1, predominantly reflecting that the receptor spends more time in long-lived closed states. In general, these results parallel those for insertions in the M3-S2 linkers (Supplemental Table 2)(Kazi *et al.*, 2014), suggesting that S1-M1 mediated processes are also driven by the D2 moving away from the membrane upon agonist binding (Figures 3C & 3D), facilitating pore opening.

### The pre-M1 helices regulate the efficiency of ion channel opening

To begin to provide mechanistic insights into the role of the β10 agonist-induced displacements in ion channel opening, we rapidly applied glutamate to single channel patches containing glycine insertions in either the GluN1 or GluN2A (Figure 6A) S1-M1s. For these experiments, we selected insertions, I546+1G in GluN1 and G542+1G in GluN2A, that were remote from membrane proximal sites where there would be more extensive local interactions. The major effect of these glycine insertions is to presumably remove the coupling between β10 and displacement of the pre-M1 helices (Twomey *et al.*, 2017).

**Figure 6.**
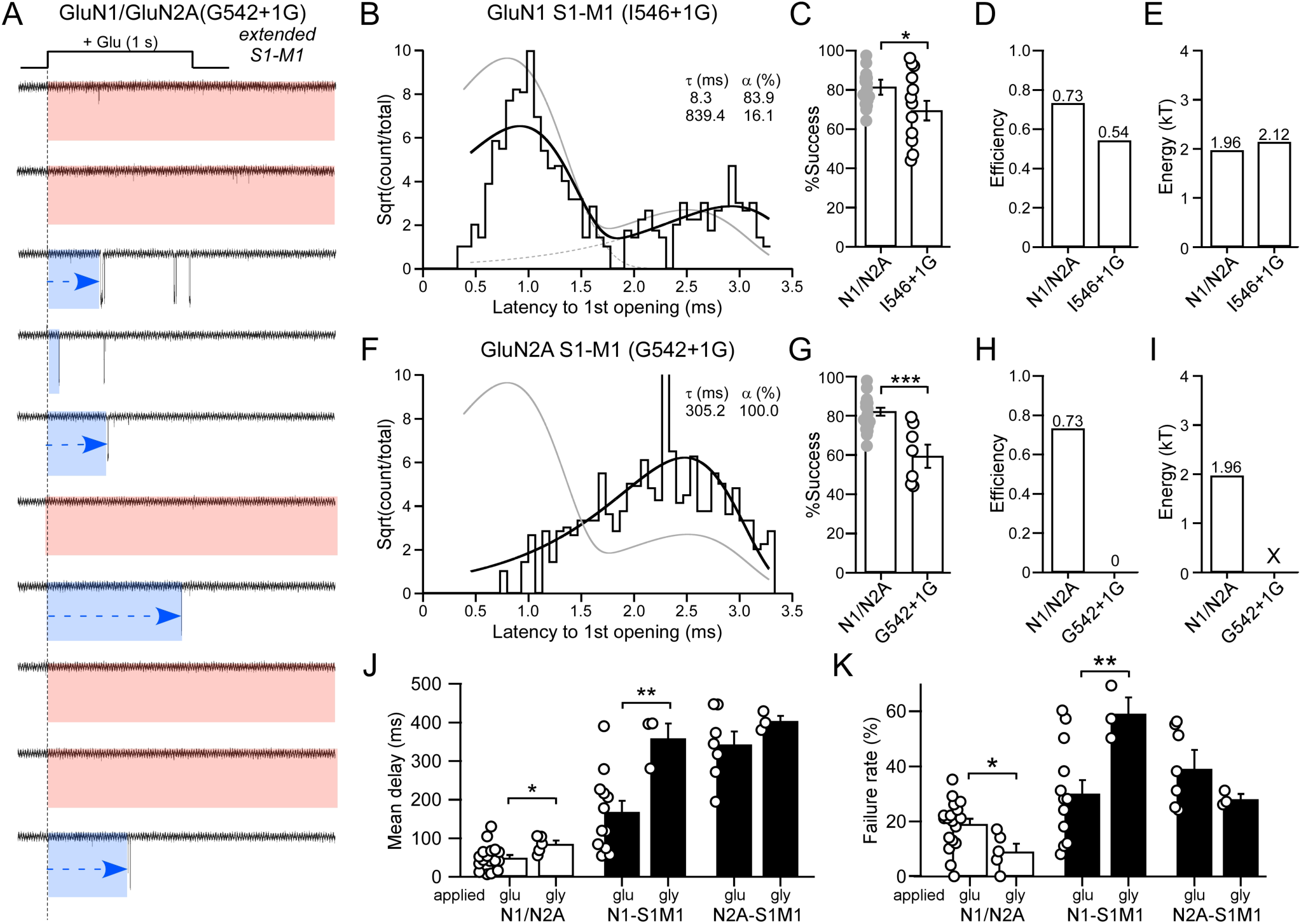
Glycine insertions in S1-M1 slow receptor activation. (**A**) Representative current traces from an outside-out single channel patch (10 consecutive traces from same recording) of a single NMDAR containing a glycine insertion in GluN2A S1-M1 (G542+1G). See Fig. 2A for details. Note numerous failures and the frequent extended delays to opening (blue). (**B-E**) Data analyzed and displayed as in Figures 4A-4D. (**B**) Dwell time histogram of the latency to 1^st^ opening for GluN1 S1-M1 (I546+1G). The histogram was best fit by 2 exponentials (dashed lines). (**C**) Success rate for the average of all patches for wild type or GluN1 S1-M1+1G (12 total patches) (global success rate, 0.65; 414 out of 639 trials). *p* < *0.05, two-tailed Student’s t-test, unpaired*. **(F-I)** Same as B-E except for a glycine insertion in the GluN2A S1-M1 (G642+1G). (**F**) Dwell time histogram was best fit by a single exponential. (**G**) Success rate for GluN2A S1-M1+1G (7 total patches) (global success rate, 0.55; 311 out of 561 trials). *p* < *0.001, two-tailed Student’s t-test, unpaired*. **(J & K)** Rapid glycine applications, in the continuous presence of glutamate, to single channel patches. Mean delay to first opening (**J**) and failures (**K**) (mean ± SEM) for wild-type and glycine insertions in GluN1 S1-M1 (I546+1G) or GluN2A S1-M1 (G542+1G). Solid bars indicate values significantly different from wild type, whereas as asterisks indicate those different from the corresponding glutamate application (*p* < *0.001, ANOVA, Tukey pairwise*). For rapid glycine applications, number of patches: Wild-type, n = 5; S1-M1: GluN1(I546+1G)/GluN2A, n = 3; and GluN1/GluN2A(G542+1G), n = 3.

For glycine insertions in the GluN1 S1-M1, the dwell time histogram was fit by 2 exponentials (Figure 6B), and there was a weak albeit significant decrease in successes (Figure 6C). Correspondingly, there was a reduction in efficiency from 0.73 to 0.54 (Figure 6D), similar to what is observed in GluN1 M3-S2 insertions (Figure 4C). However, in contrast to what is observed for GluN1 M3-S2 linker insertions, decoupling the GluN1 pre-M1 helix had limited effects on the energy required for the ‘fast’ pathway (1.96kT versus 2.12 kT) (Figure 6E). Hence, decoupling the GluN1 pre-M1 helices from agonist-induced displacements reduces the occupancy of the ‘unconstrained pre-active’ state, but does not alter the energy required for fast opening.

On the other hand, decoupling the GluN2A pre-M1 helices has a dramatic effect on the dwell time histogram, which now only has a single slow component (Figure 6F). There is a significant reduction in successes (Figure 6G). Importantly, however, given the lack of a fast component, there is no efficiency (Figure 6H). Hence, when the GluN2A pre-M1 helix is decoupled from the LBD, the receptor is no longer able to enter into the ‘unconstrained pre-active’ state.

In summary, while the on-cell data superficially suggest a similarity to what is observed when the M3 segments are decoupled from the LBD, decoupling the pre-M1 helices from the LBD has notable differences: the receptors are either inefficiently transferring energy (GluN1 pre-M1) or completely unable (GluN2A pre-M1) to enter the fast pathway. Most likely (see Discussion), entry into the fast pathway via the ‘unconstrained pre-active’ state reflects where at least one GluN2 pre-M1 is displaced and presumably at least one GluN1 pre-M1 helix, though energetically the pre-M1 helix is less critical. In contrast, the ‘constrained pre-active’ state reflects that neither GluN2 pre-M1 helices are displaced. This interpretation is consistent with recent results suggesting that the GluN2 pre-M1 helices regulate NMDAR pore opening (Gibb *et al.*, 2018; McDaniel *et al.*, 2020).

### Non-equilibrium gating for glycine binding

For our outside-out experiments, we applied glutamate in the continuous presence of glycine. To test the significance of non-equilibrium gating for glycine, we rapidly applied glycine in the continuous presence of glutamate (Figure 6J & 6K). Given the challenge of making outside-out patch recordings, we made only a limited number of these recordings and characterized the mean latency to 1st opening.

For decoupling the GluN1 pre-M1 helices, glycine activation caused the mean latency for GluN1 to be significantly slowed compared to glutamate activation (Figure 6J) as well as an increase in failure rate (Figure 6K), highlighting the role of non-equilibrium gating for the glycine binding subunit. Notably, for decoupling the GluN2A pre-M1, flipping the agonists did not have any significant effect on either the mean latency (Figure 6J) or the failure rate (Figure 6K). Hence, these experiments highlight two important points. Non-equilibrium conditions are important for defining kinetics. Second, independent of agonist, the displacement of GluN2 pre-M1 appears to always be the rate-limiting step.

### Glycine insertions in the GluN1 or GluN2A S2-M4 linkers enhance receptor gating

For S2-M4, the most proximal secondary structure in the LBD is αK in the D1 lobe (Figures 3E & 3F, 7A & 7D) (Supplemental Figures 5 & 6). In terms of glycine insertions in the S2-M4s, we followed a similar strategy as for the S1-M1s, inserting glycines (arrows) at largely equivalent positions in the GluN1 (Figure 7A) and GluN2A (Figure 7D) subunits. Insertions were also made at sites where there were numerous local interactions (red arrows) versus those regions with fewer interactions (blue arrows). In addition, there is a membrane proximal leucine (L) that in GluN2A (L812) is a disease-associated variant (Yuan *et al.*, 2014). We therefore inserted glycines around this leucine in GluN2A (Q811+1G and L812+1G) and in the homologous leucine in GluN1 (T807+1G and L808+1G).

**Figure 7.**
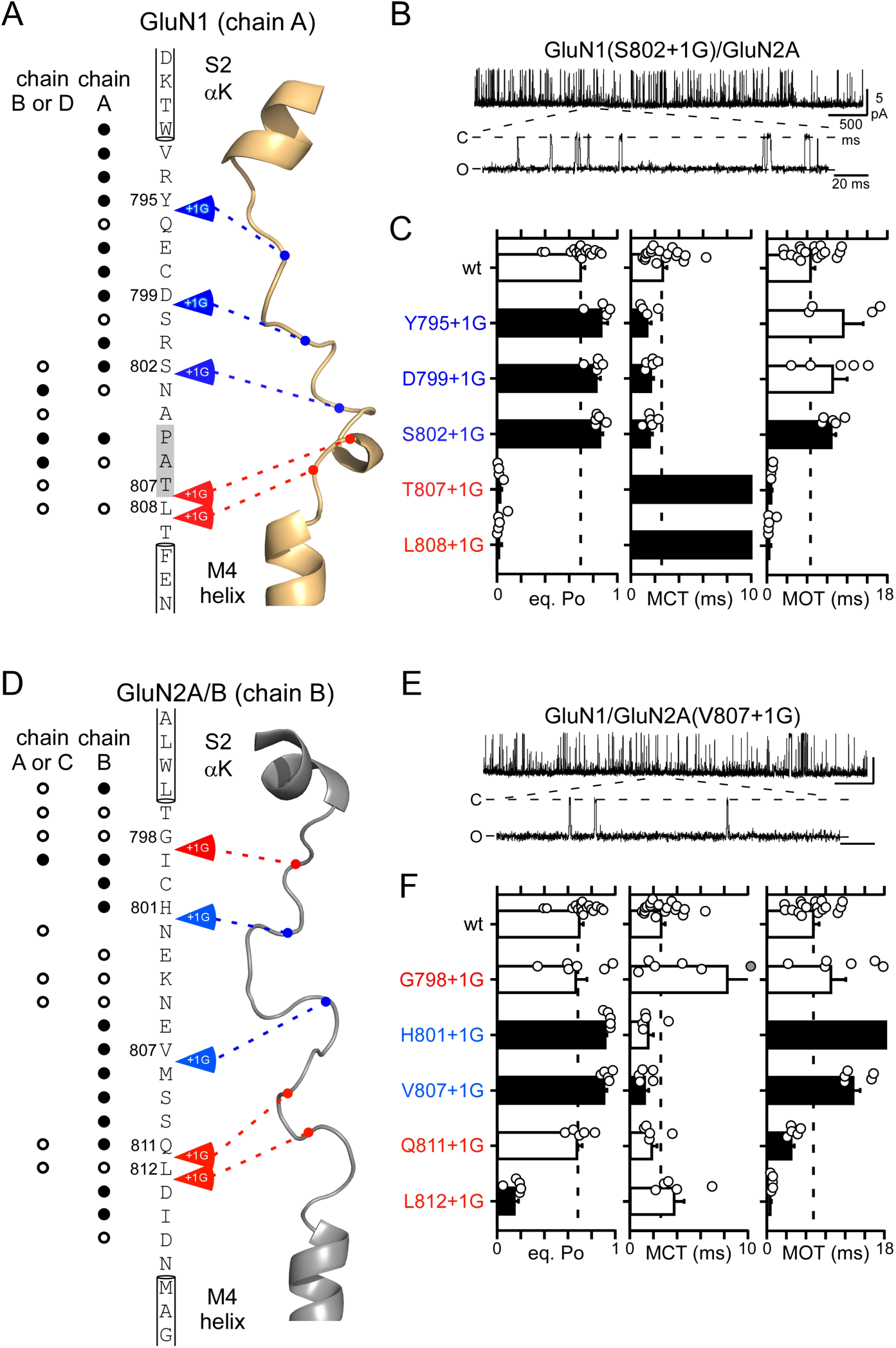
Glycine insertions in the GluN1 or GluN2A S2-M4 linkers typically potentiate NMDAR gating. **(A & D)** Primary amino acid sequences from the GluN1 and GluN2A S2-M4 linkers, as well as the last and first residues from S2 (αK) and M4 segments, respectively. Sequences displayed as in Figure 5A. Shaded bar indicates possible secondary structure. **(B & E)** Representative traces of S2-M4 glycine insertion constructs obtained using the same experimental protocols described in Figures 5B & 5E. Data are displayed as in Figures 5B & 5E. **(C & F)** Single channel *eq. P*_*o*_, MCT, and MOT (mean ± SEM) for wild-type and S2-M4 glycine insertions constructs either in GluN1 (**C**) or GluN2A (**F**). Data are shown and analyzed as in Figure 5C & 5F. MCT values for T807+1G and L808+1G and MOT values for H801+1G are incompletely shown (see Supplemental Table 4).

In contrast to insertions in the M3-S2 (Supplemental Table 2)(Kazi *et al.*, 2014) or S1-M1 (Figure 5; Supplemental Table 3) linkers, glycine insertions in the S2-M4 linkers had more diverse effects. Certain insertions had no effect on eq. Po (Figures7C & 7F, open bars), and a subset of insertions inhibited gating (T807+1G & L808+1G in GluN1 & L812+1G in GluN2A) (Figures 7C & 7F, black bars). However, these effects occurred only at insertions around the membrane-proximal leucine in GluN1 (T807+1G & L808+1G) and GluN2A (L812+1G) or at a position proximal to αK in GluN2A (G798). Notably, all of these insertions occurred at positions proximal to other linkers/transmembrane segments from other subunits (Figures 7A & 7D).

The most distinctive and consistent phenotype in the S2-M4s, especially those insertions with the least local insertions not around other residues (blue arrows), led to potentiation of gating (Figures 7B-7F). Indeed, despite a relatively high wild-type equilibrium open probability (Eq. *P*_O_) (about 0.7), glycine insertions in the S2-M4 linkers of either GluN1 (Y795+1G, D799+1G, & S802+1G) or GluN2A (H801+1G & V807+1G) subunits significantly increased eq. Po (Figures 7C & 7F). These increases in eq. Po arose most consistently through decreases in MCT in the GluN1 subunit (Figure 7C) or increases in MOT in the GluN2A subunit (Figure 7F), though these actions were not absolute.

Notably, in the S2-M4 linkers, the sites of insertion associated with potentiation were positioned between major structural elements and are not positioned around other structural elements (Figures 7A & 7D). They are also the most consistent phenotype and occur multiple times. Therefore, glycine insertions in either GluN1 or GluN2A S2-M4, which are either adding length or flexibility, enhance receptor gating.

### Glycine insertions in GluN1 and GluN2A S2-M4 improve gating efficiency

To get insights into the role of the αK and S2-M4 in agonist-induced displacements in ion channel opening, we rapidly applied glutamate to single channel patches containing glycine insertions in either the GluN1 or GluN2A (Figure 8A) S2-M4s (Figure 8). Notably, for glycine insertions in either GluN1 (Figure 8B) or GluN2A (Figure 8E), the dwell time histogram was best fit by a single exponential, and in both cases, it was only a ‘fast’ component that was comparable to what is seen in wild type. Indeed, for both constructs, failures where largely non-existent (Figures 8C & 8F), leading to nearly 100% efficiency (Figures 8D & 8G, *left panels*). Also notable was that the energy for the fast component was indistinguishable from wild type (Figures 8D & 8G, *right panels*).

**Figure 8.**
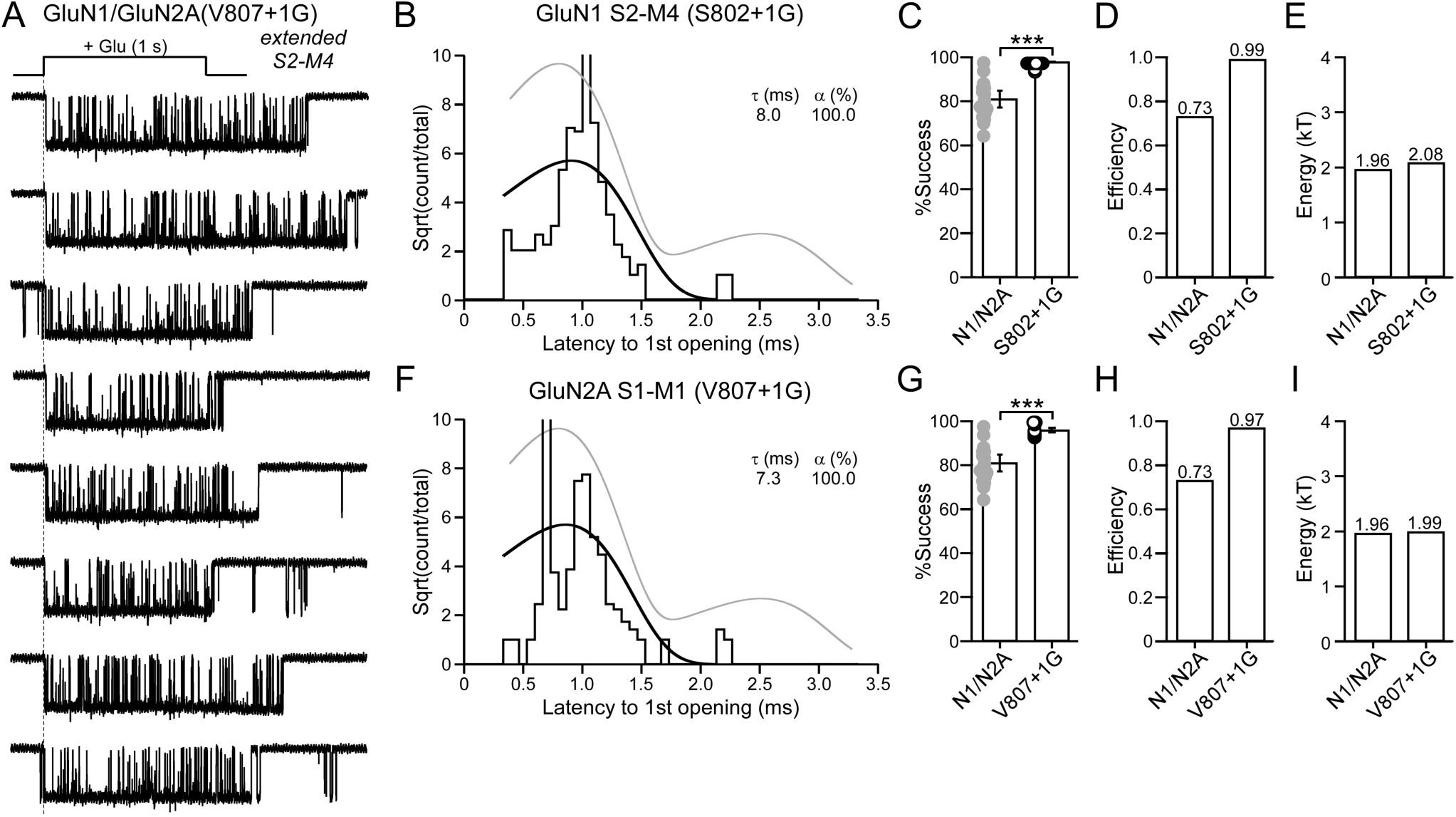
NMDARs containing glycine insertions in the GluN1 or GluN2A S2-M4 linkers favor the ‘unconstrained preactive’ state. (**A**) Representative current traces from an outside-out single channel patch (8 consecutive traces from same recording) of a single NMDAR containing a glycine insertion in the GluN2A S2-M4 (V807+1G). See Figure 2A for details. (**B-E**) Data analyzed and displayed as in Figures 4A-4D. (**B**) Dwell time histogram of the latency to 1^st^ opening for GluN1 S2-M4 (S802+1G). The histogram was best fit by a single exponential (dashed lines). (**C**) Success rate for the average of all patches for wild type or GluN1 S2-M4+1G (10 total patches) (global success rate, 0.99; 304 out of 304 trials). *p* < *0.001, two-tailed Student’s t-test, unpaired*. **(E-G)** Same as B-E except for a glycine insertion in the GluN2A S2-M4 (V807+1G). (**E**) Dwell time histogram was best fit by a single exponential. (**F**) Success rate for GluN2A S2-M4+1G (7 total patches) (global success rate, 0.97; 213 out of 219 trials). *p* < *0.001, two-tailed Student’s t-test, unpaired*. Occasionally, we observed spontaneous openings with the S2-M4 insertion: an opening occurring before application in trace 3 and possibly opening occurring long after the end of the application in traces 5, 6, and 8.

In summary, glycine insertions in either GluN1 or GluN2A S2-M4 had largely comparable effects on receptor function, dramatically enhancing receptor activation. Importantly, these S2-M4 glycine insertions did not reduce the energy of the fast transition and hence did not enhance receptor activation by easing the transition to the open state. Rather, given the near 100% efficiency, they enhance gating by allowing the receptor to always occupy the ‘unconstrained pre-active’ state. Given that the ‘unconstrained pre-active’ state reflects the displacement of the GluN2A pre-M1 helices, we assume that glycine insertions in either GluN1 or GluN2A S2-M4 facilitates displacement of the GluN2A pre-M1 helices.

## DISCUSSION

To assay the sequence of NMDAR activation, we applied glutamate to single NMDAR patches and measured the ‘latency to 1^st^ opening’, the time between when glutamate was initially applied to when the channel first opened. This latency assays events occurring between agonist binding and ion channel opening. Using this approach, we identified that wild type GluN1/GluN2A NMDARs transitions to the open state through two different intermediate pathways: a fast pathway referred to as the ‘unconstrained pre-active’ that can contribute to synaptic events, and a slow pathway referred to as the ‘constrained pre-active’ that does not (Figure 2). Given that we can make manipulations to the receptors such that it visits only the slow (Figure 6) or fast (Figure 8) pathways indicates that entry into these pathways is an either/or process and hence occurs in a parallel fashion (Figures 2H & 9).

**Figure 9.**
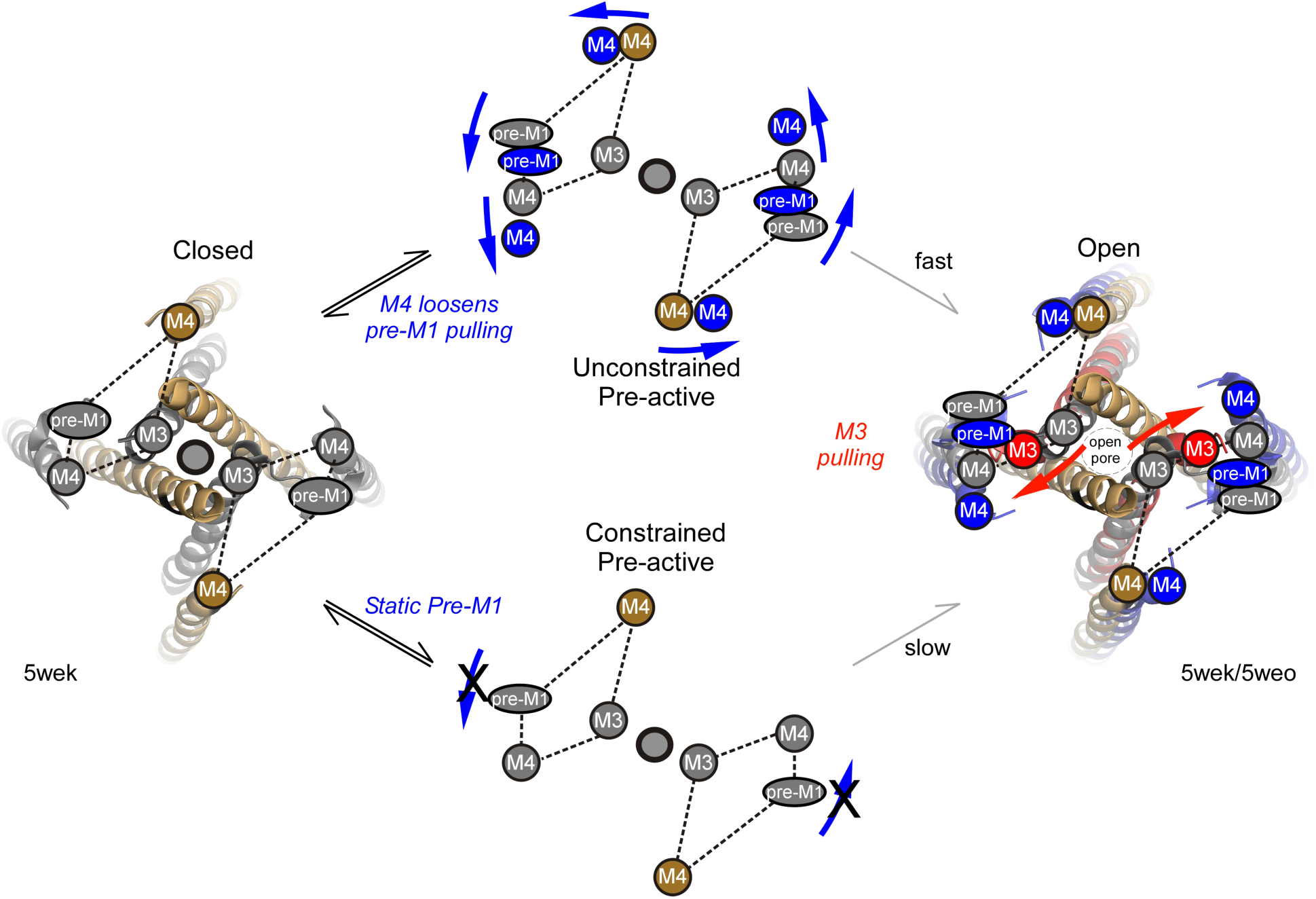
The outer structural elements must rearrange for efficient pore opening. Parallel model of NMDAR activation. Movements of the outer structural elements – GluN1 M4 (gold circle) and GluN2A pre-M1 and M4 (inner grey circles) – regulate entry from agonist-bound closed (A_4_R) (*left*) to either the unconstrained (*upper*) or constrained (*lower*) pre-active states. To efficiently enter the unconstrained state, the pre-M1 helices as well as the S2-M4/M4 must be displaced (*upper*). In contrast, if the GluN2A pre-M1 helix is not displaced, the receptor enters the constrained state, where it still can open but requires more energy to do so (*lower*). Pre-activation movements are shown in blue, whereas pore opening movements are shown in red. We show both GluN2A pre-M1 helices moving for opening, but it is more likely that only a single pre-M1 helix is required to be displaced for entry into the unconstrained preactive state (Ogden *et al.*, 2017). Stoichiometry for the S2-M4/M4s is unknown.

The entry into these intermediate pathways will determine how strongly NMDARs contribute to synaptic activity. Notably, decoupling the GluN2A pre-M1 helix from the D2-lobe displacement completely removes the ability of receptors to enter the ‘fast’ pathway (Figure 6), highlighting the importance GluN2 pre-M1 displacement to channel opening (Ogden *et al.*, 2017; McDaniel *et al.*, 2020). In addition, decoupling the GluN1 and GluN2A S2-M4, possibly by mimicking the D1-lobes downward displacement of αK (Figures 3E & 3H)(Twomey *et al.*, 2017), drives receptors into the ‘unconstrained pre-active’ (Figure 8). Thus, the outer linkers – the GluN2A pre-M1 and GluN1 and GluN2A S2-M4 – play critical roles in regulating how NMDARs contribute to synaptic signaling.

Our experiments identify that all LBD-TMD linkers – S1-M1, M3-S2, and S2-M4 – contribute to the fast process of NMDAR activation, but they do so at different points in this process. Notably, the rapid opening of the ion channel requires 1.96kT of energy (Figure 2). Not surprisingly, the energetic requirement for this fast opening step is increased when the central GluN1 (2.91kT) and GluN2 M3 (3.11kT) segments are decoupled from the LBD (Figures 4D & 4H). In contrast, decoupling the outer linkers have no effect on the energetics of the fast gating step, but rather altered the probability of entering either the unconstrained or constrained pre-active states (Figures 6 & 8). Hence, S1-M1/pre-M1 and S2-M4/M4 movements are not strongly involved in the final step of pore opening but rather determine pre-channel opening conformational changes (Figure 9).

### D1- versus D2-lobes in opening the ion channel

In terms of opening the ion channel, movements of the LBD can be divided into two categories (Figure 3)(Twomey *et al.*, 2017): D2-lobe movements, which entails β10 (S1-M1/pre-M1) and αE (M3-S2/M3), and D1-lobe movements, which entails αK (S2-M4/M4). D2-associated movements for both A/C (∼GluN1) and B/D (∼GluN2) show strong displacement away from the central axis of the mouth of the pore (Figure 3). As would be expected with mechanical coupling, decoupling pre-M1 (glycine insertions in S1-M1) or M3 (glycine insertions in M3-S2) from D2 in either GluN1 or GluN2A strongly attenuated gating (Figure 5; Supplemental Tables 2 & 3)(Kazi *et al.*, 2014). Nevertheless, as assayed by glutamate applications to outside-out single channel patches, the pre-M1 helices and the M3 segments contribute differently to the pore opening process.

Previous experiments have highlighted a gating triad between the GluN2A pre-M1 helices and the M3 segments and GluN1 S2-M4/M4s (Ogden *et al.*, 2017; Gibb *et al.*, 2018; McDaniel *et al.*, 2020). Our results strongly support the role of the GluN2A pre-M1 helix in regulating NMDAR ion channel opening. Indeed, if the GluN2A pre-M1 helices are decoupled, that is they can no longer receive sufficient mechanical displacement from the LBD, the receptor can no longer enter the unconstrained state (Figure 6F) and in terms of our efficiency parameter, no longer make any contribution to fast synaptic activity (Figure 6H).

In contrast to D2-associated linkers, the D1-associated linker show a net displacement towards the central axis of the mouth of the pore during activation (Figures 3E & 3F). Glycine insertions in either GluN1 or GluN2A S2-M4 potentiated receptor gating (Figure 6). We believe that the glycine insertions are mimicking the downward displacement of D1 towards the central axis of the mouth of the pore. It could also reflect S2-M4 unwinding (Twomey *et al.*, 2017), though we think this is unlikely given that this is fairly A/C subunit specific (Figures 3E & 3F). In either case, these results indicate that the D1 lobe is not passive in channel activation, but rather plays an active role in modulating ion channel opening. Interestingly, allosteric modulators at the extracellular ATD act by inducing a rolling movement in the LBD (Tajima *et al.*, 2016; Esmenjaud *et al.*, 2019) that may act on the αK helix, to either ‘loosen’ the S2-M4 with positive allosteric modulators or ‘tighten’ the S2-M4 with negative allosteric modulators. Hence, in addition to its active role, S2-M4 may represent a significant pathway for allosteric regulation of gating.

One surprising finding is the potentiation associated with the S2-M4 was not subunit dependent, occurring both in GluN1 and GluN2A (Figures 7 & 8). We assume that the action of S2-M4/M4 is to facilitate displacements of the GluN2A pre-M1 helices, allowing all receptors to enter into unconstrained pre-active state. Nevertheless, future experiments will be needed to fully test the specific action of the GluN1 and GluN2A S2-M4/M4 on receptor gating including the GluN2A centered gating triad. Nevertheless, if both GluN1 and GluN2A S2-M4/M4 go through the GluN2 pre-M1, then this may have to be considered a gating ‘tetrad’ rather than a gating ‘triad’ (Figure 9).

### Using latency to 1^st^ opening to assay receptor function

Assaying latency to 1^st^ opening is a powerful tool to study ion channel function (Aldrich *et al.*, 1983; Goldschen-Ohm *et al.*, 2013). We assume that the fast pathway would contribute to synaptic events whereas the slow pathway, as well as failures, would not. Hence, under our ionic conditions and because of these intermediate conformations, GluN1/GluN2A NMDARs would contribute to synaptic events about 73% of the time. For the present set of experiments, we used an external solution containing no Ca^2+^, added EDTA and at pH 8. We preferred this solution since it removed considerable uncertainty as to the effects of short- and long-term effects of Ca^2+^, Zn^2+^ and pH on receptor function (see Materials & Methods). Presumably similar conformations also exit in more physiological solutions, but the relative ratios may differ. Future experiments will be needed to address this question quantitatively.

Assaying the latency to 1^st^ opening for outside-out single channel patches has many additional uses. One intriguing use will be to flip agonists, including applying glycine or D-serine in the continuous presence of glutamate to assay how the glycine binding site (Figure 6J & 6K)(Cooper *et al.*, 2015) communicates to the TMD. In addition, more rigorous analysis including using longer agonists applications to quantify the slow component could be used to define more rigorously energetic coupling as has been done for nicotinic acetylcholine receptors (Gupta *et al.*, 2017).

### Kinetic properties of wild type NMDARs

We did not characterize rigorously the properties of the open states, in part because many of the outside-out patches had high noise. This noise issue did not impede detection of latency to 1^st^ opening, allowing us to generate large number of events, but it did make it challenging to characterize open times. Nevertheless, in those patches where we were able to reliably measure open times, there was no correlation between latency to 1^st^ opening and open time. Hence, independent of the pre-active conformation, it appears NMDAR transition to a common open state.

## Conclusion

Our experiments highlight the critical role the LBD-TMD linkers play in regulating NMDAR activity. Notable are the S2-M4/M4 segments which display numerous disease-associated variants (Yuan *et al.*, 2014; Amin *et al.*, 2018; Vyklicky *et al.*, 2018; Amin *et al.*, 2020). Further, the extracellular position of S2-M4 compared to the membrane-embedded position of M4 provides a strategic advantage for designing new drug therapies (Shi *et al.*, 2019). Importantly, knowing what role these linkers would play in the gating process allows us to consciously fine tune NMDAR gating activity.

## Acknowledgments

We thank Donna Schmidt for technical assistance; Dr. Rashek Kazi for helpful discussions and comments on the manuscript. This manuscript is dedicated to the memory of Dr. James Howe (RIP), who assisted in developing key early ideas in this manuscript. This work was supported by NIH Grants R01 NS088479 (LPW), including a minority supplement (JA).

