## Supplemental Material for "NMDA receptors require multiple pre-opening gating steps for efficient synaptic activity"

Amin and Gochman et al.

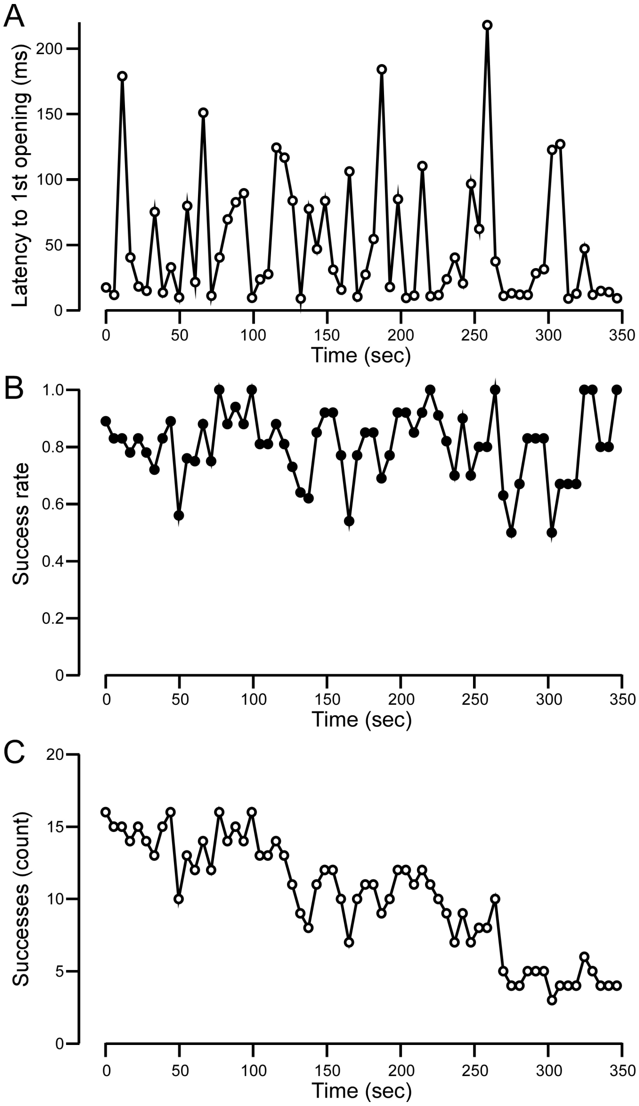

**Supplemental Figure 1. Latency to 1st opening and success rate (= 1 - failure rate) do not show time dependence (relates to Figure 2).**

Average latency to 1st opening (**A**), success rate (= 1 – failure rate) (**B**), and number of successes (**C**) for wild type GluN1/GluN2A as a function of time. A total of 18 patches were included in the analysis, ranging from 10 to 325 applications. The number of successes (**C**) decreases over time reflecting that fewer patches were available. Nevertheless, the latency to 1st opening (**A**) and success rate (**B**) showed no notable change over time.

**Supplementary Table 1. Response of wild-type GluN1/GluN2A (N1/N2A) to glutamate applications in different configurations and external solutions (relates to Figure 2).**

| | | | Act. | Deact. | Desensitization | | |
|---|---|---|---|---|---|---|---|
| **Configuration** | **External soln** | $I_{peak}$ *pA* | **10-90% rise time** *ms* | $\tau_{weighted}$ ms | $I_{peak}$ pA | *des* % | $\tau_{weighted}$ *ms* |
| | SC soln | | | | | | |
| Outside-out 1 sec | +EDTA | -160 ± 40 (16) | 4.4 ± 0.5 (13) | 70 ± 7.0 (6) | -160 ± 40 (16) | 65 ± 7 (16) | 580 ± 130 (13) |
| Whole-cell 2.5 sec | +EDTA | -2500 ± 550 (6) | 5.8 ± 0.5 (6) | 93 ± 28 (6) | -2500 ± 550 (6) | 17 ± 4* (6) | 1740 ± 370^ (6) |
| | +1 mM $CaCl_2$ | | 2 ms | | | 2.5 sec | |
| Whole-cell | +EDTA | -1550 ± 390 (5) | 7.4 ± 0.9 (4) | 44 ± 3.3 (5) | -1700 ± 390 (8) | 24 ± 5 (8) | 2420 ± 470 (8) |
| Whole-cell | -EDTA | -890 ± 220 (5) | 6.6 ± 0.8 (4) | 63 ± 7.9 (5) | -940 ± 130 (18) | 61 ± 4^ (18) | 580* ± 90 (18) |

Values shown are mean ± SEM, with the numbers in parenthesis indicating the number of patches/whole-cell recordings made for that particular parameter. **Outside-out currents**, in response to 1 sec glutamate applications, are summed currents of single channel patches (Figure 2). In some instances, typically due to a reduced number of events, too strong desensitization (difficult to measure deactivation time course), and/or too weak desensitization (difficult to measure desensitization time course), we were not able to measure all parameters for summed currents. **Whole-cell currents** were generated either using brief (2 ms) or sustained (2.5 s) glutamate applications. The rise time is the 10-90% of the rising phase of current. The decay component was fit with a double exponential with the weighted tau ($\tau_{weighted}$) shown.
***Upper two rows***. Currents were recorded in our standard single channel (SC) solution, which contained EDTA (see Materials & Methods). Tagged values are significantly less (*) or greater (^) than summed currents (*$p < 0.05$, two-tailed Student's t-test, unpaired*). Peak currents were not statistically compared.
***Lower two rows***. Currents were recorded in an external solution containing 1 mM $Ca^{2+}$ +/-EDTA (see Materials & Methods). Tagged values are significantly less (*) or greater (^) than wild type (*$p < 0.05$, two-tailed Student's t-test, unpaired*).

The extent and rate of desensitization of the summed currents was stronger and faster, respectively, than expected based on the presence of EDTA in the external solution and was more comparable to recordings done in the absence of EDTA (Supplemental Table 1). We do not know the basis for this more robust desensitization in the summed currents but it could reflect that these recordings were done in outside-out patches, and the status of the C-terminal domain, which can affect desensitization (Traynelis *et al.*, 2010; Murphy *et al.*, 2014), was altered.

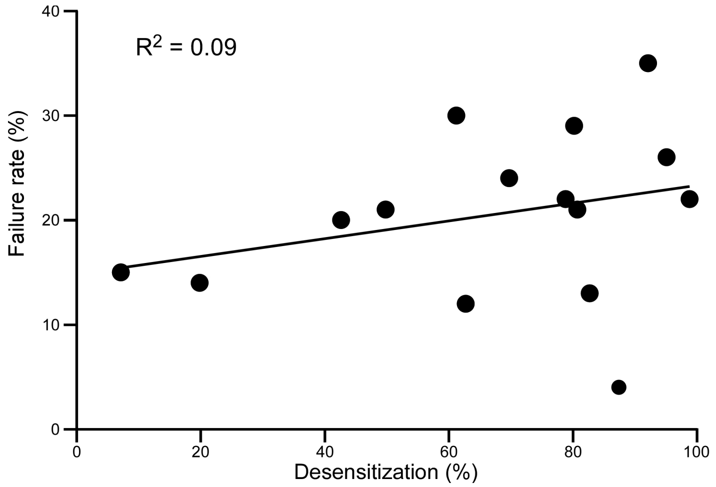

**Supplemental Figure 2. Failure rate and extent of desensitization are not correlated (relates to Figure 2).**
Failure rate plotted against the extent of desensitization for wild type GluN1/GluN2A. Only patches >10 applications were included in this analysis.

The lack of any correlation between failure rates and the extent of desensitization suggest that desensitization does not contribute significantly to failure rates.

**Supplemental Table 2. Single channel properties of glycine insertions in the GluN1 or GluN2A M3-S2 (Kazi et al., 2014) (relates to Figure 4).**

| Construct | eq. $P_{open}$ | MCT *ms* | MOT *ms* |
|---|---|---|---|
| N1/N2A | 0.67 ± 0.06 | 4.3 ± 0.8 | 9.0 ± 0.9 |
| N1(G666+1G)/N2A (=N1(G648+1G)[3]) | 0.36 ± 0.06* | 10 ± 1.5^ | 4.9 ± 0.6* |
| N1(I664+1G)/N2A | 0.29 ± 0.03* | 7.3 ± 0.8^ | 2.7 ± 0.2* |
| N1(E662+1G)/N2A | 0.27 ± 0.05* | 9.9 ± 1.4^ | 3.1 ± 0.4* |
| N1/N2A(G664+1G) = N2A(G645+1G) | 0.08 ± 0.02* | 86.8 ± 30^ | 4.5 ± 0.4* |
| N1/N2A(D660+1G) | 0.01 ± 0.009* | 510 ± 310 | 2.4 ± 0.5* |
| N1/N2A(F658+1G) | 0.02 ± 0.007* | 260 ± 160 | 2.3 ± 0.1* |
| N1/N2A(E656+1G) | 0.003 ± 0.001* | 470 ± 140 | 1.2 ± 0.2* |

Values are taken from Supplemental Table 1 from (Kazi *et al.*, 2014). They are only shown as a reference.
Tagged values are significantly less (*) or greater (^) than wild type (*$p < 0.05$, two-tailed Student's t-test, unpaired*).

Note in Kazi et al. (2014) we used the numbering for the mature protein (the numbering shown here minus the signal peptide: GluN1 (18); GluN2A (19)).

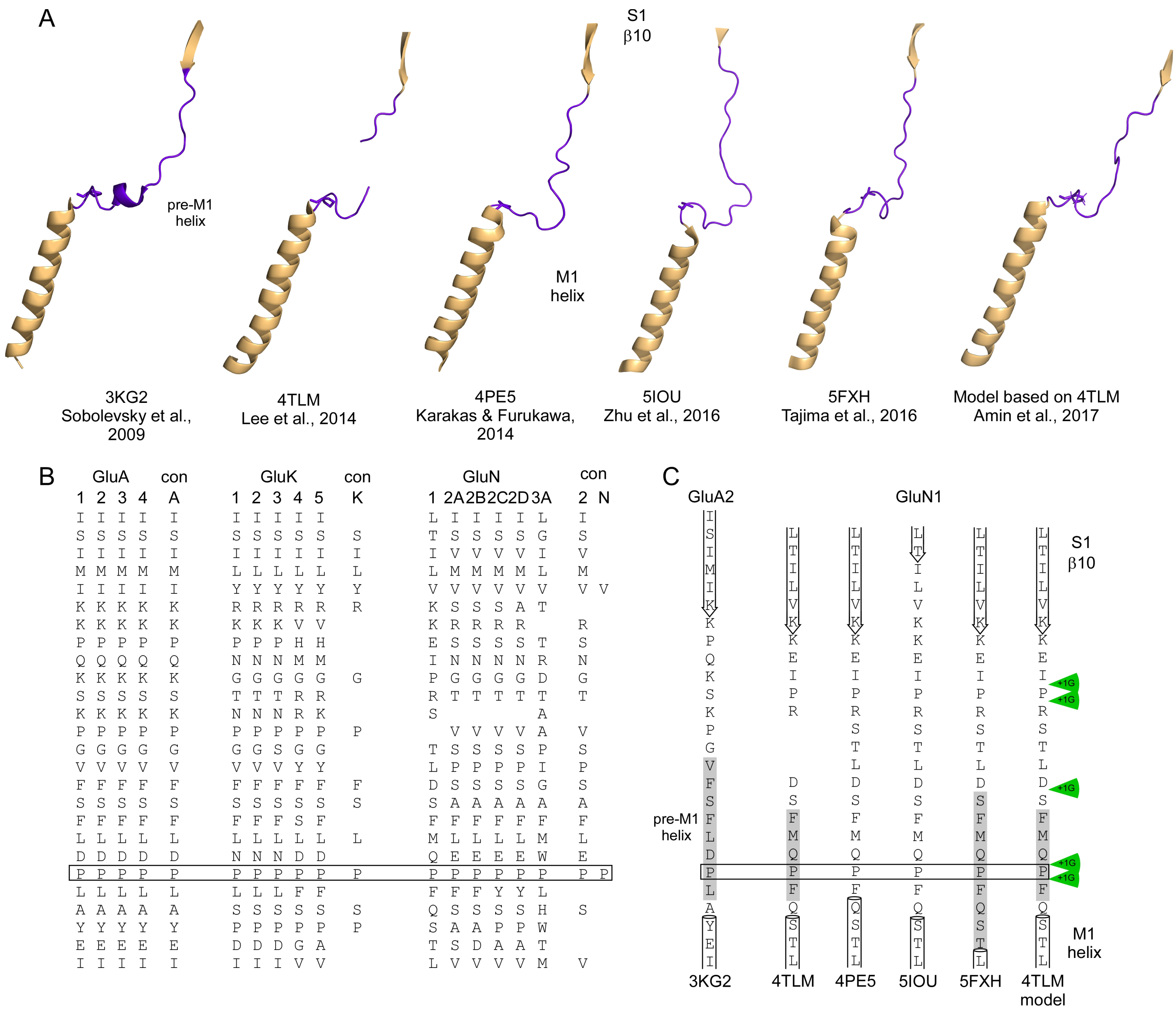

**Supplemental Figure 3. Topology and sequence alignment of S1-M1 linkers and adjacent regions in the AMPAR GluA2 and NMDAR GluN1 subunit (relates to Figure 5).**

**(A)** Comparison of high-resolution structures and a computational model based on 4TLM (Amin *et al.*, 2017) of S1-M1 in AMPAR (GluA2; 3KG2, subunit A) (Sobolevsky *et al.*, 2009) and NMDAR GluN1 (GluN1/GluN2B; 4TLM (Lee *et al.*, 2014), 4PE5 (Karakas & Furukawa, 2014), 5IOU (Zhu *et al.*, 2016), & 5FXH (Tajima *et al.*, 2016), computational model). The last secondary structural element in S1 is β10. In the different structures, the end of β10 and the start of the M1 helix varies, with the specific amino acid transitions shown in (C). Notable in the S1-M1 linker is a pre-M1 helix. Secondary structure elements comprising the LBD (β10) and TMD (M1 helix) are colored light orange while the S1-M1 linkers are highlighted in purple. Also in purple is the side group of a highly conserved proline residue in S1-M1 (see B) that was used to align structures.

**(B)** Sequence alignment of iGluR subtypes (AMPAR (GluA2), kainate (GluK), & NMDAR (GluN)) encompassing S1-M1 and adjacent regions. Conserved (con) residues are shown to the right of alignments. For NMDAR subunits, the conserved residues are shown for GluN2 (2) or for all NMDAR (N) subunits except for GluN3B. A proline (boxed) is the only completely conserved residue in this region.

**(C)** Primary amino acid sequences of the S1-M1 and adjacent regions in the different structures shown in A. Location of secondary structural elements are indicated by downward arrows (β10), gray box (pre-M1 helix), or cylinders (M1 helix). The locations and transitions of secondary structures were taken from the DSSP image of the PDB. Secondary structures were indicated when more than 2 positions showed such a structure. Residues not shown were not resolved in the structures. Green arrows indicate where glycines were inserted.

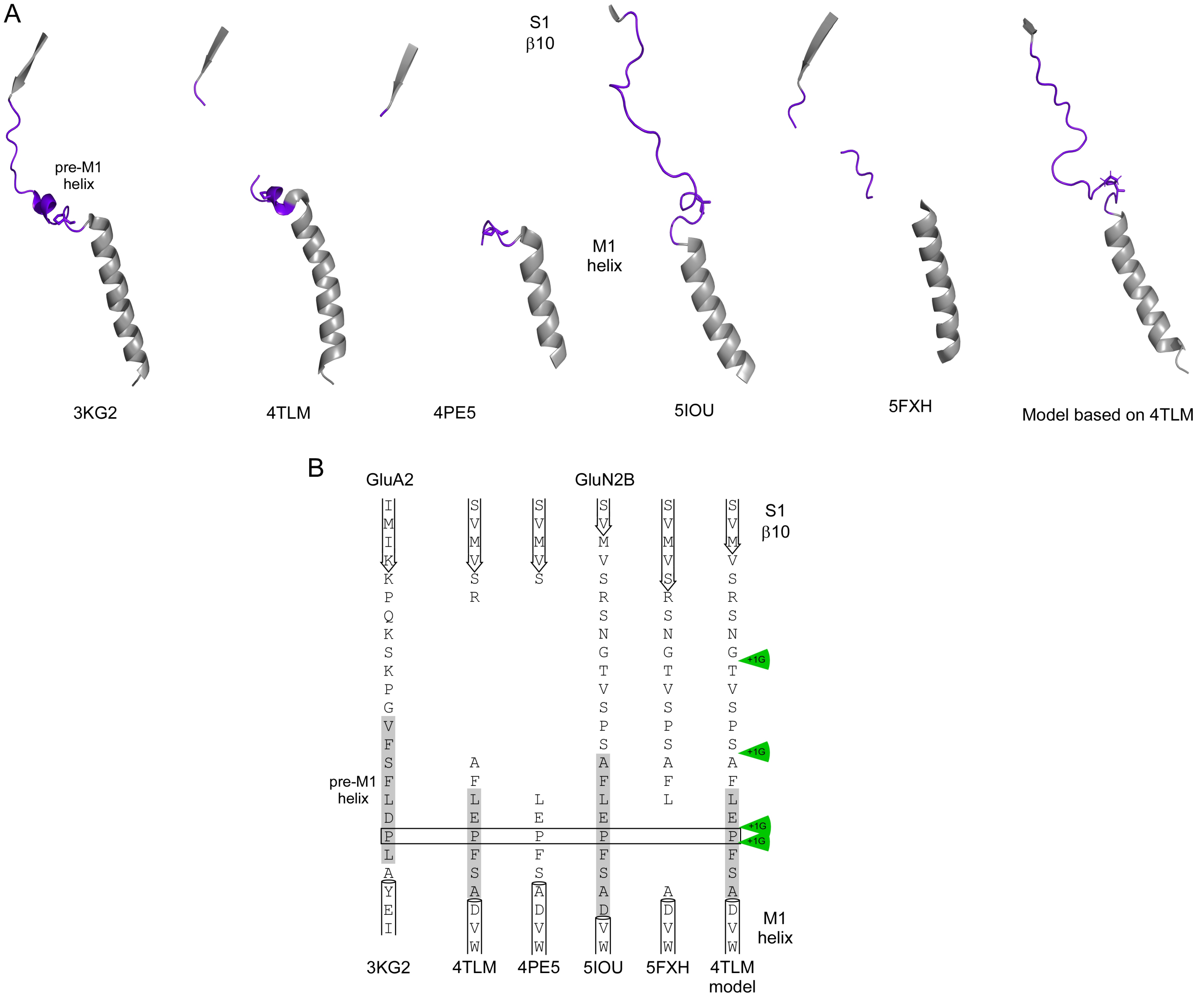

**Supplemental Figure 4. Topology and sequence alignment of S1-M1 linkers and adjacent regions in the AMPAR GluA2 and NMDAR GluN2B subunit (relates to Figure 5).**

**(A)** Comparison of high-resolution structures, as in Figure S3A, except encompassing GluA2 (subunit B) and GluN2B S1-M1. The last secondary structure in GluN2B S1 is β10. As for GluN1 (Figure S1), the end of β10 and the start of the M1 helix varies in the different structures, with the specific amino acid transitions shown in (B). Secondary structure elements comprising the LBD (β10) and TMD (M1 helix) are colored gray while the S1-M1 linkers are highlighted in purple. The conserved proline used for sequence alignment is in purple.

**(B)** Primary amino acid sequences of the S1-M1 and adjacent regions in the different structures shown in (A). Locations of secondary structural elements are indicated by downward arrows, cylinders or gray bars. Green arrows indicate where glycines were inserted.

**Supplemental Table 3. Single channel properties of wild-type GluN1/GluN2A (N1/N2A) or single glycine insertions in GluN1 or GluN2A S1-M1 (relates to Figure 5).**

| Construct | Total events (# of patches) | *i* *pA* | eq. $P_{open}$ | MCT *ms* | MOT *ms* |
|---|---|---|---|---|---|
| N1/N2A | 1,512,458 (18) | -6.9 ± 0.1 | 0.70 ± 0.03 | 2.7 ± 0.3 | 6.6 ± 0.6 |
| N1(I546+1G)/N2A | 301,397 (4) | -7.5 ± 0.4 | 0.47 ± 0.06* | 4.7 ± 1.0 | 3.9 ± 0.3* |
| N1(P547+1G)/N2A | 963,084 (5) | -7.1 ± 0.2 | 0.68 ± 0.05 | 2.1 ± 0.3 | 4.5 ± 0.5* |
| N1(D552+1G)/N2A | 1,104,164 (5) | -5.7 ± 0.2* | 0.43 ± 0.08* | 2.0 ± 0.5 | 1.3 ± 0.2* |
| N1(Q556+1G)/N2A | 37,580 (4) | -7.7 ± 0.9 | 0.05 ± 0.03* | 27.1 ± 6.4^ | 0.8 ± 0.2* |
| N1(P557+1G)/N2A | 492,157 (4) | -7.2 ± 0.3 | 0.36 ± 0.04* | 6.3 ± 1.0^ | 3.5 ± 0.6* |
| N1/N2A(G542+1G) | 65,983 (4) | -7.4 ± 0.7 | 0.06 ± 0.01* | 73.1 ± 7.6^ | 3.9 ± 0.4* |
| N1/N2A(S547+1G) | 11,284 (5) | -8.0 ± 0.8 | 0.01 ± 0.005* | 170 ± 50^ | 1.2 ± 0.1* |
| N1/N2A(E551+1G) | 38,814 (4) | -7.9 ± 0.4 | 0.06 ± 0.01* | 61.3 ± 7.2^ | 3.6 ± 0.2* |
| N1/N2A(P552+1G) | *nd* | | | | |

Values shown are mean ± S.E.M. for single-channel current amplitude (*i*), equilibrium open probability (eq. $P_o$), mean closed time (MCT), and mean open time (MOT). Single channel currents were recorded in the on-cell mode at approximately -100 mV and analyzed in QuB (see Materials & Methods). Numbers of patches are in parentheses to the right of total events. Eq. $P_o$ is the fractional occupancy of the open states in the entire single-channel recording, including long lived closed states.
Tagged values are significantly less (*) or greater (^) than those of wild type (*$p < 0.05$, two-tailed Student's t-test, unpaired*).
*nd*, no glutamate-activated currents were detected in the on-cell single channel patches for N1/N2A(P552+1G). This construct was not tested in the whole-cell mode but lack of surface expression is not uncommon for mutations at N2A(P552)(Ogden *et al.*, 2017).

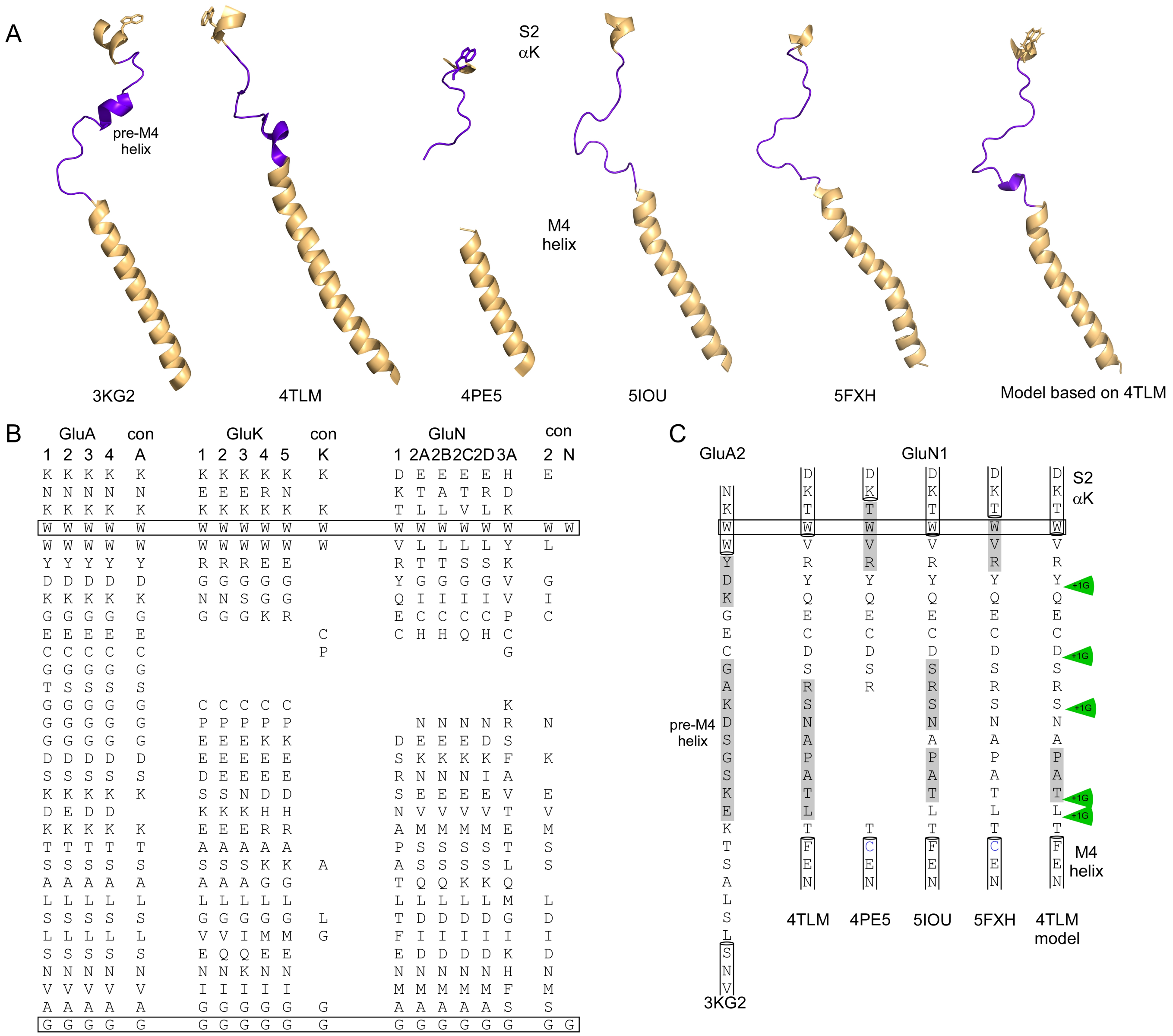

**Supplemental Figure 5. Topology and sequence alignment of S2-M4 linkers and adjacent regions in the AMPAR GluA2 and NMDAR GluN1 subunit (relates to Figure 7).**

**(A)** Comparison of high-resolution structures, as in Figure S3A, except for GluA2 and GluN1 S2-M4. The last secondary structural element in S2 is αK. Present in the AMPAR S2-M4 linker is a pre-M4 helix. Secondary structure elements comprising the LBD (αK) and TMD (M4 helix) are colored light orange while the S2-M4 linkers are highlighted in purple. A highly conserved tryptophan residue (see B), used for sequence alignment, is shown in either light orange (3KG2, 4TLM, 5IOU, 5FXH, model) or purple (4PE5).

**(B)** Sequence alignment, as in Figure S3B, shown for S2-M4 linkers and adjacent regions for the iGluR subtypes. The only completely conserved residues in this region are a tryptophan (W) and a glycine (G) in the upper M4 (boxed). Conserved (con) residues are shown to the right of alignments.

**(C)** Primary amino acid sequences of the S2-M4 and adjacent regions in the different structures shown in (A). The locations and transitions of secondary structures were taken from the DSSP image of the PDB. Locations of secondary structural elements are indicated by cylinders. Residues not shown were not included in the structures. Green arrows indicate where glycines were inserted.

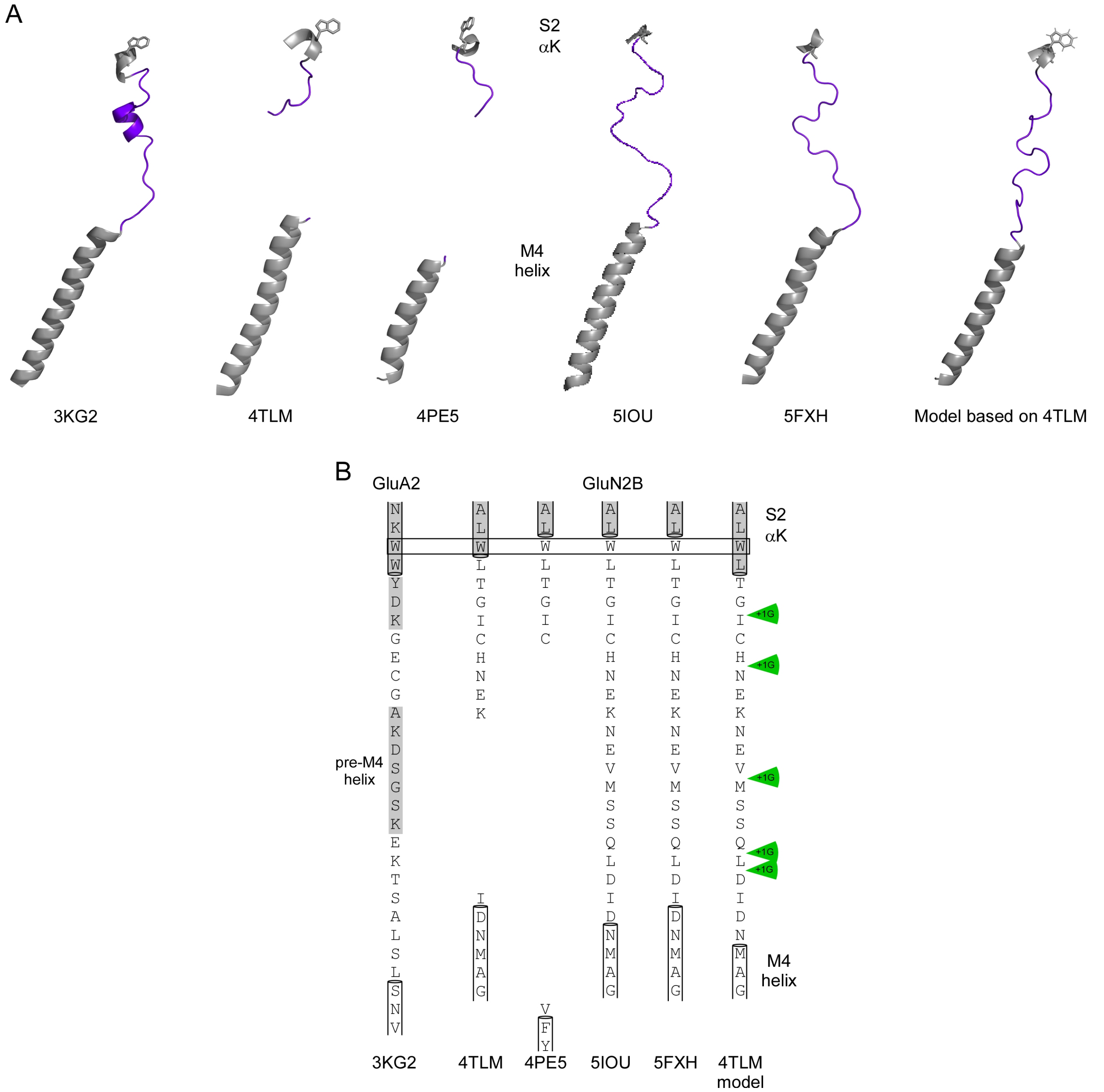

**Supplemental Figure 6. Topology and sequence alignment of S2-M4 linkers and adjacent regions in the AMPAR GluA2 and NMDAR GluN2B subunit (relates to Figure 7).**

**(A)** Comparison of high-resolution structures, as in Figure S3A, except encompassing GluA2 and GluN2B S2-M4. As shown in Figure S5A, the end of αK and the start of the M4 helix varies, with the specific amino acid transitions shown in (B). The AMPAR pre-M4 helix is not clearly present in GluN2B S2-M4. Secondary structure elements comprising the LBD (αK) and TMD (M4 helix) are colored gray while the S2-M4 linkers are highlighted in purple. The side group of a highly conserved tryptophan residue in S2 (see Figure S5B) used for sequence alignment is shown in gray.

**(B)** Primary amino acid sequences of the S2-M4 and associated regions in the different structures shown in A. Location of secondary structural elements are indicated by cylinders. Residues not shown were not included in the structures. Green arrows indicate where glycines were inserted.

**Supplemental Table 4. Single channel properties of glycine insertions in GluN1 or GluN2A S2-M4 (relates to Figure 7).**

| **Construct** | **Total events (# of patches)** | ***i*** ***pA*** | **eq. $P_{open}$** | **MCT** ***ms*** | **MOT** ***ms*** |
|---|---|---|---|---|---|
| N1/N2A | 1,512,458 (18) | -6.9 ± 0.1 | 0.70 ± 0.03 | 2.7 ± 0.3 | 6.6 ± 0.6 |
| N1(Y795+1G)/N2A | 86,605 (4) | -7.0 ± 0.2 | 0.87 ± 0.05^ | 1.4 ± 0.3* | 11.5 ± 2.9 |
| N1(D799+1G)/N2A | 102,676 (5) | -5.7 ± 0.4 | 0.83 ± 0.03^ | 1.7 ± 0.2* | 9.9 ± 2.1 |
| N1(S802+1G)/N2A | 303311 (5) | -7.3 ± 0.1 | 0.86 ± 0.02^ | 1.6 ± 0.2 | 9.7 ± 0.6^ |
| N1(T807+1G)/N2A | 57,692 (4) | -7.7 ± 0.8 | 0.03 ± 0.01* | 37.1 ± 14.9 | 0.7 ± 0.1* |
| N1(L808M+1G)/N2A | 59,063 (5) | -6.3 ± 0.6 | 0.02 ± 0.02* | 30.2 ± 9.3^ | 0.4 ± 0.2* |
| N1/N2A(G798+1G) | 427,214 (6) | -6.9 ± 0.4 | 0.67 ± 0.09 | 8.3 ± 5.4 | 11.0 ± 2.2 |
| N1/N2A(H801+1G) | 111,130 (5) | -6.6 ± 0.3 | 0.92 ± 0.01^ | 1.6 ± 0.4 | 20.5 ± 4.6^ |
| N1/N2A(V807+1G) | 125,552 (5) | -6.7 ± 0.3 | 0.91 ± 0.02^ | 1.3 ± 0.3* | 13.3 ± 1.0^ |
| N1/N2A(Q811+1G) | 424,137 (5) | -4.8 ± 0.7* | 0.68 ± 0.04 | 1.9 ± 0.4 | 3.8 ± 0.4* |
| N1/N2A(L812+1G) | 226,981 (5) | -4.8 ± 0.3* | 0.15 ± 0.03* | 3.8 ± 0.8 | 0.6 ± 0.1* |

Values shown are analyzed as in Supplemental Table 5.
Tagged values are significantly less (*) or greater (^) than wild type (*$p < 0.05$, two-tailed Student's t-test, unpaired*).

**References used in Supplemental Material**

Amin JB, Salussolia CL, Chan K, Regan MC, Dai J, Zhou HX, Furukawa H, Bowen ME & Wollmuth LP. (2017). Divergent roles of a peripheral transmembrane segment in AMPA and NMDA receptors. *J Gen Physiol* **149,** 661-680.

Karakas E & Furukawa H. (2014). Crystal structure of a heterotetrameric NMDA receptor ion channel. *Science* **344,** 992-997.

Kazi R, Dai J, Sweeney C, Zhou HX & Wollmuth LP. (2014). Mechanical coupling maintains the fidelity of NMDA receptor-mediated currents. *Nat Neurosci* **17,** 914-922.

Lee CH, Lu W, Michel JC, Goehring A, Du J, Song X & Gouaux E. (2014). NMDA receptor structures reveal subunit arrangement and pore architecture. *Nature* **511,** 191-197.

Murphy JA, Stein IS, Lau CG, Peixoto RT, Aman TK, Kaneko N, Aromolaran K, Saulnier JL, Popescu GK, Sabatini BL, et al. (2014). Phosphorylation of Ser1166 on GluN2B by PKA is critical to synaptic NMDA receptor function and Ca2+ signaling in spines. *J Neurosci* **34,** 869-879.

Ogden KK, Chen W, Swanger SA, McDaniel MJ, Fan LZ, Hu C, Tankovic A, Kusumoto H, Kosobucki GJ, Schulien AJ, et al. (2017). Molecular Mechanism of Disease-Associated Mutations in the Pre-M1 Helix of NMDA Receptors and Potential Rescue Pharmacology. *PLoS Genet* **13,** e1006536.

Sobolevsky AI, Rosconi MP & Gouaux E. (2009). X-ray structure, symmetry and mechanism of an AMPA-subtype glutamate receptor. *Nature* **462,** 745-756.

Tajima N, Karakas E, Grant T, Simorowski N, Diaz-Avalos R, Grigorieff N & Furukawa H. (2016). Activation of NMDA receptors and the mechanism of inhibition by ifenprodil. *Nature* **534,** 63-68.

Traynelis SF, Wollmuth LP, McBain CJ, Menniti FS, Vance KM, Ogden KK, Hansen KB, Yuan H, Myers SJ & Dingledine R. (2010). Glutamate receptor ion channels: structure, regulation, and function. *Pharmacol Rev* **62,** 405-496.

Zhu S, Stein RA, Yoshioka C, Lee CH, Goehring A, McHaourab HS & Gouaux E. (2016). Mechanism of NMDA Receptor Inhibition and Activation. *Cell* **165,** 704-714.
